# The impact of lure, trap type, and Fluon application on the capture of longhorned beetles (Coleoptera: Cerambycidae) in the subtropical forests of southeastern Louisiana, USA

**DOI:** 10.64898/2026.08.18.745610

**Authors:** Chiranjivi Sharma, Winter Sheline, Elise Grossman, Nicole Jean-Williams, Jorge Macias-Samano, Robert R. Setter, Todd D. Johnson

**Affiliations:** Department of Entomology, Louisiana State University, Baton Rouge, LA, USA; Synergy Semiochemicals Corp., Delta, British Columbia, Canada

**Keywords:** Semiochemical, trap design, forest insect monitoring, attractant, pheromone

## Abstract

The expansion of global trade has increased the risk of unintentional introductions of woodboring insects outside their native ranges, which can cause considerable ecological and economic damage. Monitoring efforts for native and non-native species of woodborers rely heavily on semiochemical-baited traps. With the goal of improving detection efforts of longhorned beetles (Coleoptera: Cerambycidae) and their associates, we conducted field bioassays in loblolly pine dominated and bottomland hardwood forest ecosystems, which are representative of subtropical Louisiana. We tested the impact of attractant composition, trap design, and the application of a friction reducing compound on the capture of longhorned beetles. We compared the Synergy Multi-Trap System with and without Fluon, and the Synergy Funnel Trap II with Fluon, baited with multicomponent cerambycid lures and ethanol or ethanol alone. In total, we collected 5,303 beetles comprising 44 species. Traps baited with cerambycid lures and ethanol captured ∼11X more individuals than the traps baited with ethanol. We identified novel attractants for 6 species of longhorned beetles. When Fluon was applied, both the Synergy Multi-Trap System and Funnel Trap II were equally effective at capturing cerambycids. Application of Fluon significantly increased the trap capture of the Synergy Multi-Trap System by ∼4X compared to the same trap type without Fluon. Our results contribute to our understanding of the factors influencing trap captures and the chemical ecology of woodboring insects thereby helping the development of early detection and rapid response monitoring programs in subtropical regions of the United States and elsewhere.

## Introduction

Increasing globalization has accelerated unintentional introductions of non-native woodboring insects through international trade, particularly via infested wood and wood products (Dobbs and Brodel, 2004; Levine & D’Antonio, 2003; Roques et al., 2010; Haack et al., 2014). Because woodborer larvae develop endophytically within woody tissues, infestations are difficult to detect during routine visual inspections at ports of entry (Greenwood et al., 2023). Consequently, many non-native woodborers have been introduced outside of their native ranges, threatening woody plants in managed and unmanaged forests globally (Brockerhoff et al., 2006; Nowak et al., 2001).

In forest ecosystems, woodboring beetles in the family Cerambycidae, commonly known as the longhorned beetles, play critical roles as nutrient recyclers, by initiating and hastening the decomposition of stressed, weakened or dead woody materials (Duffy, 1953; Hanks, 1999; Lingafelter, 2007; Linsley, 1959), often in association with saproxylic fungi (Haack, 2017; Parker et al., 2006; Stokland et al., 2012; Ulyshen, 2016). In their native habitats, most cerambycids colonize damaged or weakened woody hosts, while a smaller fraction of species attack healthy trees and herbaceous plants (Craighead, 1923; Slansky, 1987; Solomon, 1995; Hanks, 1999). When introduced into new environments, however, longhorned beetles can cause substantial ecological and economic damage due to the absence of co-evolved natural enemies (Keane & Crawley, 2002), reduced biotic resistance (Elton, 1958), genetic adaptability, and favorable host and environmental conditions (Maron & Vilà, 2001). For instance, losses resulting from the Asian longhorned beetle, *Anoplophora glabripennis* Motschulsky (Cerambycidae: Lamiinae), first detected in the United States in 1996, have likely exceeded projected economic losses of $1.23 trillion dollars (adjusted for inflation) in the United States (Nowak et al., 2001; US Inflation Calculator, 2025).

Although regulations such as the International Standards for Phytosanitary Measures No. 15 (IPSM 15) have reduced the risk of new introductions (Allen et al., 2017; Haack, 2006), the cross-border movement of woodboring longhorned beetles persists (Haack et al., 2014), with increasing detections of non-native species far beyond their native ranges (Eyre & Haack, 2017; Lawson et al., 2018). As such, effective monitoring tools are essential for the early detection of this ecologically and economically important group of woodboring insects. Because many species of woodboring longhorned beetles use long-range volatile pheromones to locate mates and hosts (Hanks & Millar, 2016), semiochemical-baited traps are often used in monitoring programs for non-native species (Allison & Redak, 2017). These traps exploit hardwired responses to reproductive signals to improve detection efficiency, and support efforts to prevent introductions while safeguarding terrestrial ecosystems.

Over the past decade, identification of cerambycid pheromones has greatly expanded due to the discovery of high levels of semiochemical parsimony across the Cerambycidae, wherein identical or similar pheromone components are often shared among closely related species at the genus, tribe, and subfamily levels of taxonomic organization (Hanks & Millar, 2016). This finding has led to the development of the strategy known as pheromone identification by proxy (Millar et al., 2019), wherein deployment of pheromones of one species of cerambycid can be used to simultaneously attract, detect, and validate the attractive response of multiple species of cerambycids within regions (Miller et al., 2017; Ray et al., 2015; Sweeney et al., 2014; Wickham et al., 2014), across regions (Hanks et al., 2018), and even between continents (Miller et al., 2017; Fan et al., 2019). Such results demonstrate that advances in the detection and management of invasive cerambycids can be made by studying the chemical ecology of relatively benign species of cerambycids in their native habitats.

The detection of woodboring cerambycids can be further enhanced by combining multiple pheromone components, as well as host-volatiles in multi-lure blends (Collignon et al., 2016; Fan et al., 2019; Flaherty et al., 2019; Hanks et al., 2012; Hanks & Millar, 2018; Molander & Larsson, 2018; Rassati et al., 2019; Wong et al., 2012). For example, an 8-component pheromone blend captured 376 cerambycid species representing 8 subfamilies and 60 tribes across Asia, North America, the Caribbean, and Australia (Roques et al., 2023). Because it is challenging to predict which species are likely to arrive, and at what frequency/number (i.e., their propagule pressure), multi-component pheromone blends provide effective monitoring tools for both targeted and untargeted cerambycid surveillance programs. These programs enable detection of known species at low population densities as well as phylogenetically related invaders whose pheromones are known or yet to be discovered (Hanks et al., 2012; Roques et al., 2023; Wong et al., 2012).

While cerambycid pheromonal lures are critical for attracting beetles to the vicinity of a trap, its design can strongly influence its capture efficiency. At short ranges, visual cues are often used by cerambycids to assess the presence of suitable hosts (Bernays & Chapman, 1994; Lyu et al., 2015; Otálora-Luna et al., 2013) or mates (Johnson et al., 2019) prior to landing. After cerambycids have made the decision to land, trap features such as available surface area, surface area orientation (i.e., funnel vs. panel), and surface texture can all impact the likelihood of detecting target and non-target species of cerambycids. For instance, Allison et al. (2014) compared three different types of traps that varied in their design (e.g., funnel vs. panel) and size and found that that multiple funnel traps captured significantly more beetles than panel and stovepipe traps. Findings may vary though according to region and target species; in a similar study, Graham et al. (2012) found that panel traps captured 1.5X more cerambycid beetles than funnel traps.

To further enhance trap efficiency and beetle retention, trap surfaces are often treated with a fluoropolymer-dispersion (Fluon), which increases surface slipperiness and reduces the ability of beetles to alight or walk on trap surfaces, thereby increasing likelihood that they fall into the collection jar (Graham et al., 2010; Graham & Poland, 2012). However, recent U.S. and European Union regulations restricting per- and polyfluoroalkyl substances (PFAS), including fluoropolymers, raise concerns about future availability of Fluon (Environmental Protection Agency, 2020; European Chemicals Agency, 2024). Additionally, Fluon is costly and requires repeated applications because its effectiveness decreases over time (Dong et al., 2023). Although a single application cost approximately $4.50 per panel trap (Graham & Poland, 2012), expenses ($6.50 if adjusted for inflation; US Inflation Calculator, 2025) rapidly accumulate in large-scale monitoring and Early Detection and Rapid Response (EDRR) programs that deploy thousands of traps. Therefore, developing traps with materials that eliminate the need for repeated Fluon application could provide a more economical and efficient alternative for regulatory agencies, forest managers, and researchers, which we are evaluating using Synergy Multi-Trap Systems.

Reliable and cost-effective traps are essential for EDRR monitoring and eradication programs targeting non-native cerambycids across diverse forest ecosystems. To address these management needs, we conducted a six-month field experiment across two sites in subtropical, southeastern Louisiana, to determine if a new plastic formulation would increase capture efficiency and reduce the need for Fluon application on traps. This was tested with the Synergy Multi-Trap System (hereafter, MTS) with and without Fluon treatment. Our findings were compared to the standard Funnel Trap II (hereafter, FT-II) treated with Fluon. As part of our study, we used six-component pheromone lures paired with low-release ethanol bags to attract a wide-diversity of cerambycids to test our hypothesis. Because there is little published information regarding the cerambycid fauna of subtropical habitats, including southeastern Louisiana, we included control traps baited with ethanol alone to identify new information on attractants by cerambycids in our study. This experiment was designed to maximize capture efficiency and accurately identify all cerambycid species present, reflecting the priorities of researchers and regulatory agencies concerned with biodiversity surveillance and invasion risk assessment.

## Materials and methods

### Site of study

Our field experiment was carried out at two distinct sites in southeastern Louisiana. The first site was the Bob R. Jones Idlewild Research Station in Clinton, LA (30.801 N, −90.956 W; Fig. 1A, B), a 526-hectare loblolly pine (Pinaceae: *Pinus taeda* L.) dominated forest with hardwood bottomland species including, musclewood (*Carpinus caroliniana* Walter), American beech (*Fagus grandifolia* Ehrh.), oak (*Quercus* spp. L.), and black cherry (*Prunus serotina* Ehrh.). Our second site was the Audubon Nature Center in New Orleans, LA (29.924 N, −90.131 W; Fig. 1A, C), a 35-hectare bottomland hardwood ecosystem characterized by bald cypress (*Taxodium distichum* (L.) Rich.), water oak (*Quercus nigra* L.), mulberry (*Morus* spp. L.), rough-leaf dogwood (*Cornus drummondii* Mey.), red maple (*Acer rubrum* L.), and willow (*Salix* spp. L.).

**Figure 1.**
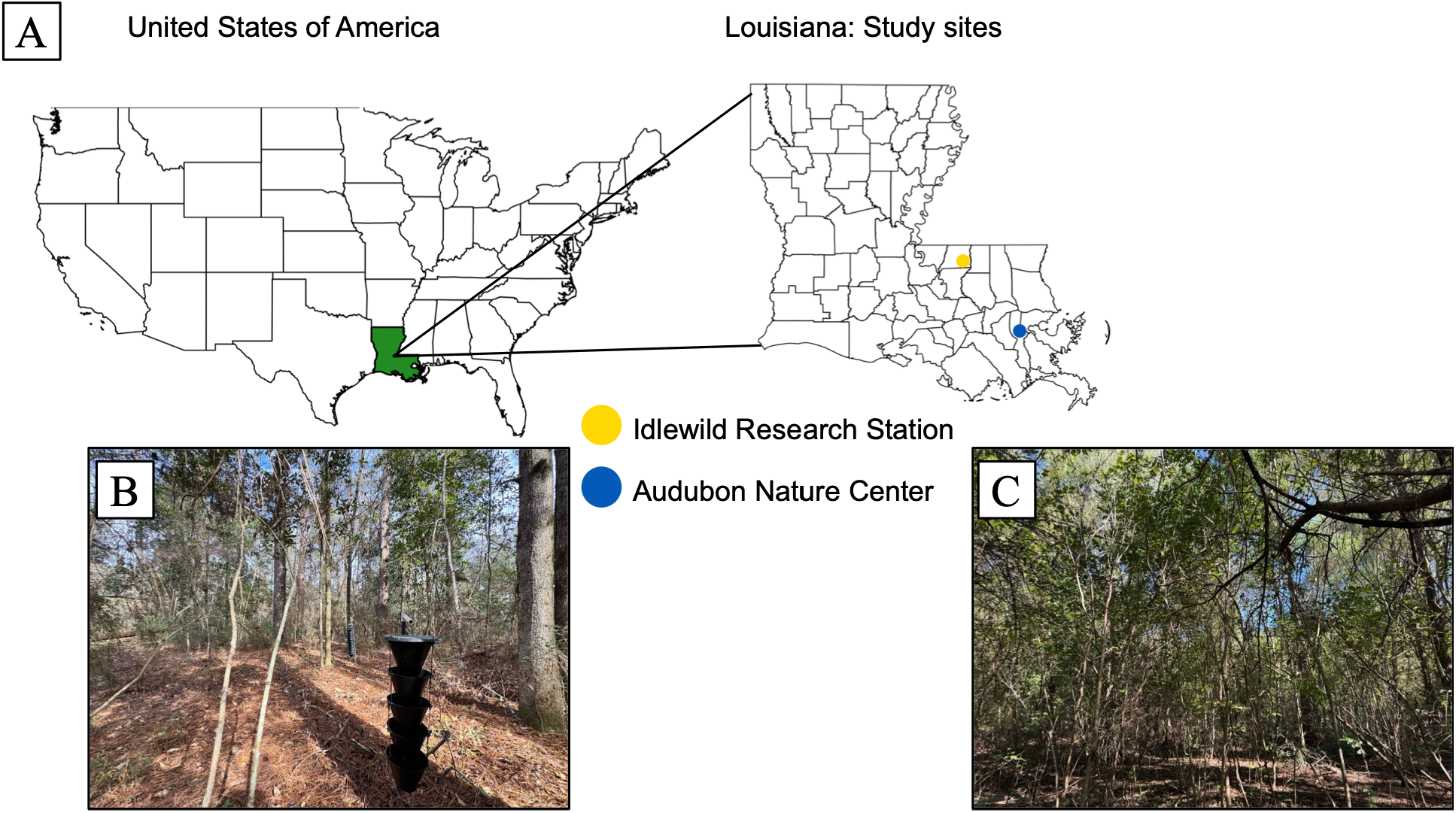
Study sites: (A) Geographic location of the Idlewild Research Station and Audubon Nature Center; (B, C) representative habitat views of each site, respectively, in southeastern subtropical Louisiana, USA.

### Source of lures

All pheromone lures were acquired from Synergy Semiochemicals Corp. (Delta, BC, Canada). Traps were baited with a blend of cerambycid pheromones (Synergy product #3620) having an average release rate of 4.5 mg/day at 25 °C and low-release ethanol lure (Synergy product #3344; hereafter, ethanol) with an average release rate of 40 mg/day at 25 °C. The cerambycid pheromone blend contained six semiochemicals: (*E*)-6,10-dimethylundeca-5,9-dien-2-ol (fuscumol), (*E*)-6,10-dimethylundeca-5,9-dien-2-yl acetate (fuscumol acetate), (5*E*)-6,10-dimethylundeca-5,9-dien-2-one (geranylacetone), a racemic mixture of 3-hydroxy-2-hexanone (3,2-ketol), 2-methylbutanol, and *anti*-2,3-hexanediol (*anti*-C6-diol) (Supp Fig. S1). An ethanol lure was used either alone as control, or in combination with the cerambycid pheromone blend. The composition and loading rates of each constituent in the cerambycid pheromone blend are detailed in Table 1.

**Table 1.** Composition and loading rate of chemical constituents of 2g cerambycid pheromone blend bubble dispenser.

| Pheromone | Mixture % | Load (2g bubble) |
| --- | --- | --- |
| Fuscumol | 18 | 0.36 |
| Fuscumol acetate | 19 | 0.36 |
| Geranyl acetone | 9 | 0.18 |
| 3-hydroxy-2-hexanone | 18 | 0.36 |
| 2-methylbutanol | 18 | 0.36 |
| *Anti*-2,3-hexanediol | 18 | 0.36 |

### Experimental trap design and treatment structure

A total of 36 traps comprising 12 Synergy Multi-Trap System (MTS) treated with a fluoropolymer-dispersion (Fluon PTFE; Synergy EZ Fluon DIY Kit, Synergy Semiochemicals Corp., Delta, BC, Canada; Fig. 2A), 12 MTS without Fluon application (Fig. 2B), and 12 Synergy Funnel Trap II (FT-II) with Fluon application (Fig. 2C) procured from Synergy Semiochemicals Corp. (Delta, BC, Canada) were deployed for the experiment. At each site, 18 traps representing all experimental treatments (defined by the combination of lure composition, Fluon application, and trap type) were deployed in a linear transect: 6 MTS with Fluon application, 6 MTS without Fluon application, and 6 FT-II with Fluon application. These traps were selected because they are widely standardized in North American forest pest surveys, align with the methodological consistency with USDA Forest Service monitoring programs and have proven efficiency in intercepting the longhorned beetles (Allison et al., 2014). The Synergy MTS and FT-II primarily differ in interception surface area and funnel geometry. The MTS has 5 vertical funnels made of different plastic with a hardener intended to make the surface smoother, having a steeper angle of inclination, broader at the top (r = 11.1 cm), and twice the height (17.1 cm) of those in FT-II comprising 11 vertical funnels that are narrower at the top (r = 9.35 cm) with a vertical height (8.5 cm). These differences alter the trap’s plume structure (i.e., aerodynamics) and visual profiles, potentially affecting beetle flight interception and landing behavior, thereby allowing us to test how trap design (MTS vs FT-II both treated with Fluon) influences capture rates.

**Figure 2.**
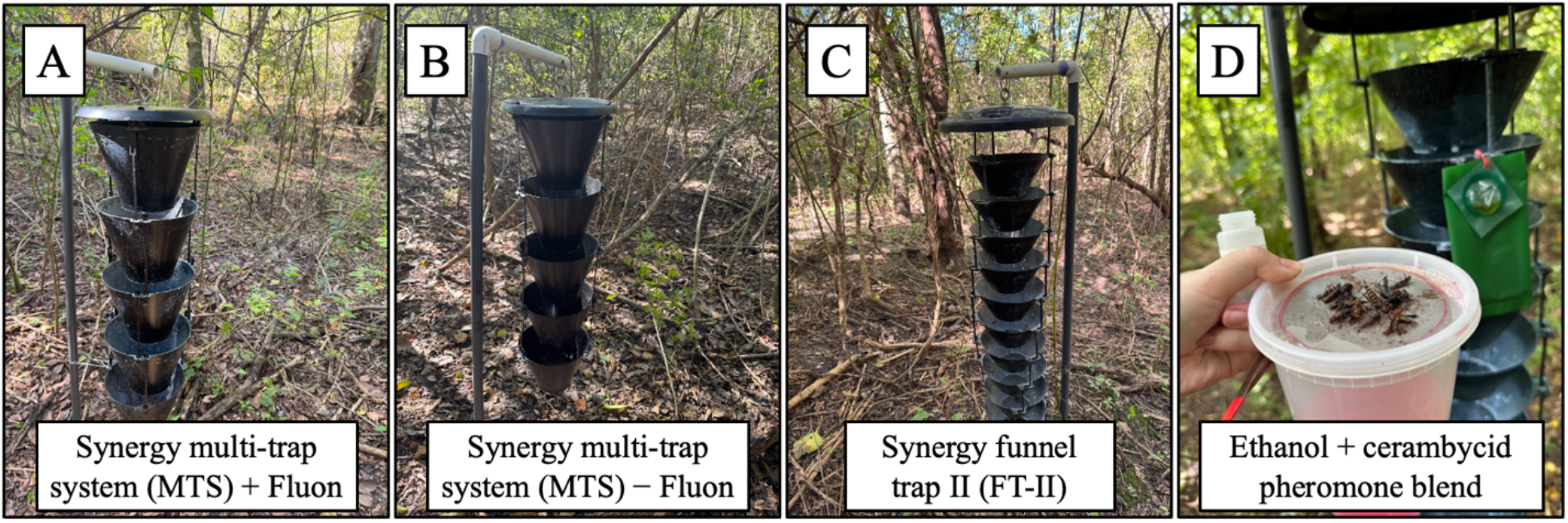
Synergy Multi-trap System (MTS) (A) with, and (B) without Fluon application, and (C) Synergy Funnel Trap II (FT-II) with Fluon set in a linear transect with (D) lure suspended laterally on the edge of the traps.

To test our hypothesis that the material used to manufacture MTS would be sufficiently effective to eliminate the need for Fluon application, 6 MTS treated with Fluon and 6 MTS untreated with Fluon were deployed at each site. To test whether lure composition impacts the attraction of longhorned beetles to traps, 5 traps of each type were baited with a cerambycid pheromone blend paired with an ethanol lure (Fig. 2D), and 1 trap of each type was baited with low release ethanol lure alone as controls. We paired cerambycid pheromones blend and ethanol lures because alcohols, including ethanol, are emitted by stressed, weakened or damaged host trees, which are preferred host condition of many woodborers, and enhance attraction of longhorned beetles to traps (Hanks et al., 2012; Miller & Sweeney, 2025; Rice et al., 2024). Because ethanol itself is not attractive to many species of longhorned beetles (Sweeney et al., 2004), ethanol lures were used as controls to assess the impact of lures on the collection of beetles. We did not use blank control traps to avoid the problem of zero catches across all replicates, which could lead to zero variance and make statistical analysis difficult (Reeve & Strom, 2004).

### Trapping deployment and duration of experiment

Prior to trap setup, we randomized the starting position of all traps within our linear transects. Inverted L-shaped frames constructed from black colored polyvinyl chloride (PVC) pipe were used for suspending traps by mounting those frames on 1-meter-long steel reinforcing bar partially driven and stabilized in the ground (Graham et al., 2010). The top of each trap was positioned approximately 1.5 meters, while the bottom of the trap was about 0.5 meters above the ground. Lures were suspended laterally on the edge of each trap (Fig. 2D). Traps were separated from each other by at least 5 meters, a distance shown to minimize interference between treatments when traps are deployed in linear transects in the field (Wong et al., 2017). Similarly, Miller and Crowe (2018) have shown no effect of trap spacing on capturing ambrosia and bark beetles, weevils, longhorn beetles, bark beetle predators and associates when pheromone baited traps were spaced by 6 or 12 m compared to 2 m. Each trap basin had a wet collection cup of 1L capacity filled approximately half-way with a non-toxic propylene glycol solution (Inhibited Propylene Glycol 95%, ChemWorld, Kennesaw, GA) to kill and preserve captured beetles.

The experiment was conducted from 11 July through 21 November 2024, for a total of 36 trap check dates. Traps were checked twice a week, and their positions were rotated along the linear transect during each trap check. This rotation ensured that position effects that each trap may experience (i.e., proximity to stressed trees or placement at the interior vs edge of the forest) were controlled throughout our study. The captured longhorned beetles were identified to species level, and taxonomic names and authorities were verified using Lingafelter (2007), Bezark (2019) and Slipinski and Escalona (2016), in combination with comparisons to curated reference specimens housed in the Louisiana State Arthropod Museum.

### Data preparation

To evaluate our hypotheses, we generated three subsets of data (Suppl. Fig. 2). The first subset contained the trap catch of 20 species of cerambycids that were found to be significantly affected by at least one treatment factor in our study (Table 2). We used this dataset to test the hypothesis (H_1_) that lure would impact response by cerambycids (Suppl. Fig. 2, Step 3). Our second subset of data contained counts of these beetles collected only from MTS and FT-II traps with Fluon; these data were used to test the hypothesis (H_2_) that trap type would influence the collection of cerambycids (Suppl. Fig. 2, Step 4A). Lastly, our third subset contained counts of beetles collected only from MTS traps with and without Fluon; these data were used to test the hypothesis (H_3_) that Fluon would impact capture of cerambycids (Suppl. Fig. 2, Step 4B).

**Table 2.** Cerambycid species that were used for statistical analysis with their estimate, SE and p-value of each treatment factor.

| Species | Estimate ± SE | | | p-value | | |
| --- | --- | --- | --- | --- | --- | --- |
|  | Trap type | Fluon | Lure | Trap type | Fluon | Lure |
| *Aegomorphus modestus* (Gyllenhal) | | 1.35 ± 0.52 | | | <0.01 | |
| *Aegomorphus quadrigibbus* (Say) | | 1.2 ± 0.78 | 2.93 ± 0.74 | | <0.01 | <0.01 |
| *Atrypanius haldemani* (LeConte) | | 2.66 ± 0.74 | 2.80 ± 0.60 | | <0.01 | <0.01 |
| *Curius dentatus* (Newman) | 0.40 ± 0.21 | 1.61 ± 0.28 | 4.59 ± 0.72 | 0.04 | <0.01 | <0.01 |
| *Elaphidion mucronatum* (Say) | | 2.62 ± 0.44 | 3.04 ± 0.36 | | <0.01 | <0.01 |
| *Graphisurus fasciatus* (DeGeer) | -0.47 ± 0.21 | 1.31 ± 0.36 | 3.01 ± 0.46 | 0.03 | <0.01 | <0.01 |
| *Graphisurus triangulifer* (Haldeman) | | | 3.67 ± 1.02 | | | <0.01 |
| *Leptostylopsis argentatus* (du Val) | -0.60 ± 0.30 | 2.11 ± 0.63 | 1.95 ± 0.38 | 0.04 | <0.01 | <0.01 |
| *Leptostylopsis planidorsus* (LeConte) | | | 2.46 ± 0.76 | | | <0.01 |
| *Leptostylus transversus* (Gyllenhal) | | 2.30 ± 1.05 | 1.61 ± 0.63 | | 0.03 | 0.01 |
| *Lepturges angulatus* (LeConte) | | 2.25 ± 0.46 | 3.50 ± 0.53 | | <0.01 | <0.01 |
| *Liopinus alpha* (Say) | | 1.79 ± 0.76 | | | 0.02 | |

| Species | Trap type | Estimate ± SE | | Trap type | p-value | |
| --- | --- | --- | --- | --- | --- | --- |
|  |  | Fluon | Lure |  | Fluon | Lure |
| *Neoclytus acuminatus* (Fabricius) | | | 0.94 ± 0.41 | | | 0.02 |
| *Neoclytus mucronatus* (Fabricius) | | 2.08 ± 0.22 | 4.74 ± 0.50 | | <0.01 | <0.01 |
| *Neoclytus scutellaris* (Olivier) | | | 3.11 ± 1.03 | | | <0.01 |
| *Orthosoma brunneum* (Forster) | | | 1.10 ± 0.37 | | | <0.01 |
| *Plectromerus dentipes* (Olivier) | | 2.25 ± 0.74 | 3.76 ± 1.01 | | <0.01 | <0.01 |
| *Styloleptus biustus* (LeConte) | | 1.76 ± 0.14 | 3.14 ± 0.13 | <0.01 | <0.01 | <0.01 |
| *Styloleptus biustus minuens* (LeConte) | | 1.74 ± 0.41 | | | <0.01 | |
| *Xylotrechus colonus* (Fabricius) | | 1.81 ± 0.39 | 2.06 ± 0.29 | | <0.01 | <0.01 |

Our experimental design with more treatments than controls for lure hypothesis was to maximize the collection of both number of individuals and species of longhorned beetles. Prior to analysis, this difference in number of traps with or without the cerambycid lure was standardized by calculating the number of beetles captured per trap/treatment/date for each site (Suppl. Fig. 2, Step 2). Additionally, for each species of cerambycid, we removed trap data on check dates with no capture records of beetles (Suppl. Fig. 2, Step 1). Lack of capture in all traps on a date may have represented conditions when beetles were not in flight due to inclement weather, or trap dates outside of their phenological window, neither of which were conducive to evaluating our hypotheses (Johnson et al., 2021; Rice et al., 2020).

### Data analysis

Binary count data was analyzed using statistical software R version 4.4.1 (R Core Team, 2024). Prior to analysis, diagnostic plots were used to determine the structure of the data, including residual versus fitted data and Q-Q plots in R, as well as the relationship between mean and variance of each species of beetle across treatments (Zeileis et al., 2008; Zuur et al., 2009). We found that our data were not normally distributed and over dispersed (variance > mean) and therefore were best modeled using a negative binomial error distribution fitted with a log-link function (Bolker et al., 2009).

For each species of cerambycid collected in our study, we ran full interactive and additive generalized linear mixed models (GLMMs) with the glmmTMB package (Brooks et al., 2017). Models were constructed with treatments (lure type, trap type, and Fluon application) as fixed effects while trap-check date and site as random effects. Model comparisons were also made for significance between additive vs interactive models. The best models were selected based on the lowest Akaike Information Criterion (AIC) and highest Bayesian Information Criterion (BIC) values (Supp. Table S1). Trap-check dates and sites were modeled as random effects to account for repeated measurements of same traps over time, and for the nesting of traps within sites, thereby addressing temporal and spatial non-independence and minimizing the risk of pseudo-replication and inflated Type I error (Bolker et al., 2009; Harrison et al., 2018; Zuur et al., 2009).

Our best models were additive (trap type + Fluon application + lure type + 1/site + 1/trap check date; Supp. Table S1). The model summaries indicated that 20 species had significant responses to at least one treatment factor, with no interactive effects found (Table 2). Each of these 20 species of beetles was captured on at least 4 trap-check dates and represented by at least 18 specimens (Supp. Table S2). The capture records of these 20 species of beetles were pooled/aggregated to present our main treatment effects.

Tukey’s Honestly Significant Difference (HSD) post hoc test (α = 0.05) was used to conduct pairwise comparison among treatment means only when the models identified significant treatment effects. Although trap-check dates were significant for several species, we are not reporting it here, as variation in trap captures across dates likely reflect fluctuations in weather conditions and other biotic and abiotic factors that were not specifically evaluated as part of our study (Hanks et al., 2018).

## Results

### Cerambycids captured across trap types and forest habitats

Between July and November 2024, we captured a total of 5,303 longhorned beetles, comprising 44 species across the subfamilies of Cerambycinae (n = 13), Lamiinae (n = 28), and Prioninae (n = 2). We also collected one species of distenid, which was previously included in the Cerambycidae as the subfamily Disteniinae (Švácha and Lawrence, 2014; Table 3). Of the total species captured, 36 species (n = 2918) were captured in MTS while 40 species (n = 2385) were captured in FT-II. *Elaphidion tectum* LeConte, *Knulliana cincta cincta* Drury, *Smodicum cucujiforme* Say, and *Astylopsis fascipennis* Schiefer were only captured in MTS while *Distenia undata* Fabricius, *Mallodon dasytomus* Say, *Enaphalodes atomarius* Drury, *Parelaphidion aspersum* Haldeman, *Aegomorphus morrisii* Uhler, *Hyperplatys aspersa* Say, and *Pogonocherus mixtus* Haldeman were captured only in FT-II (Table 3).

**Table 3.**
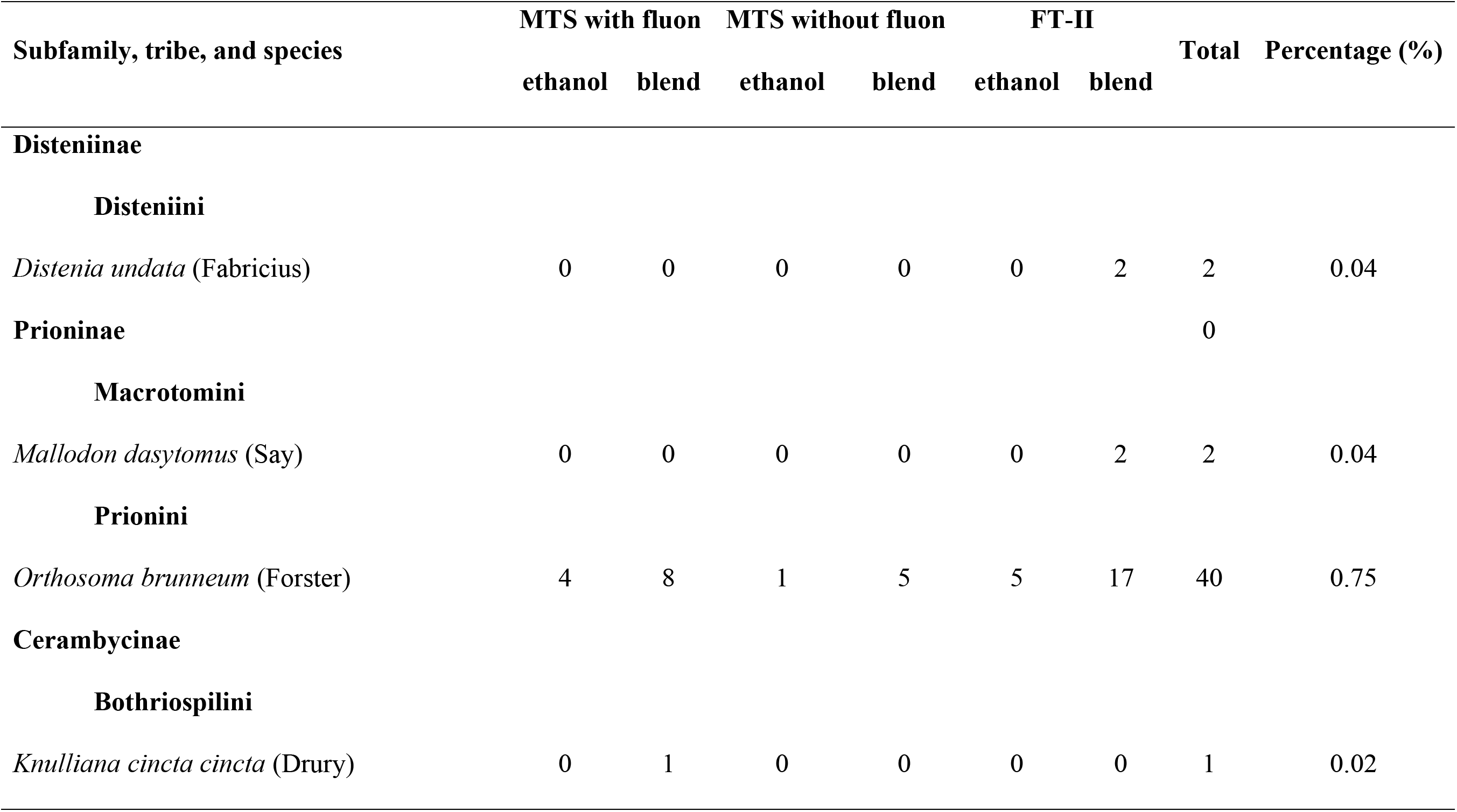
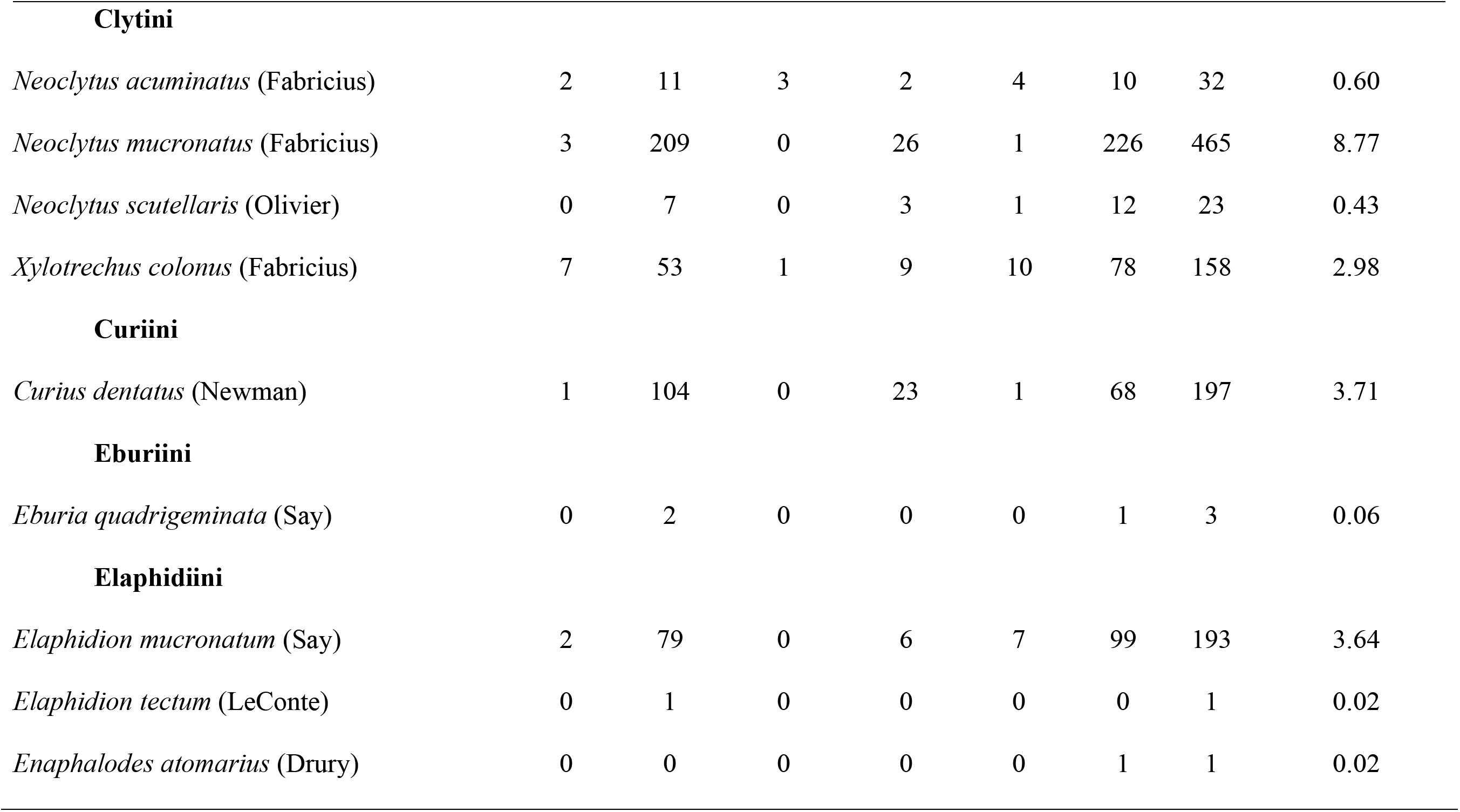
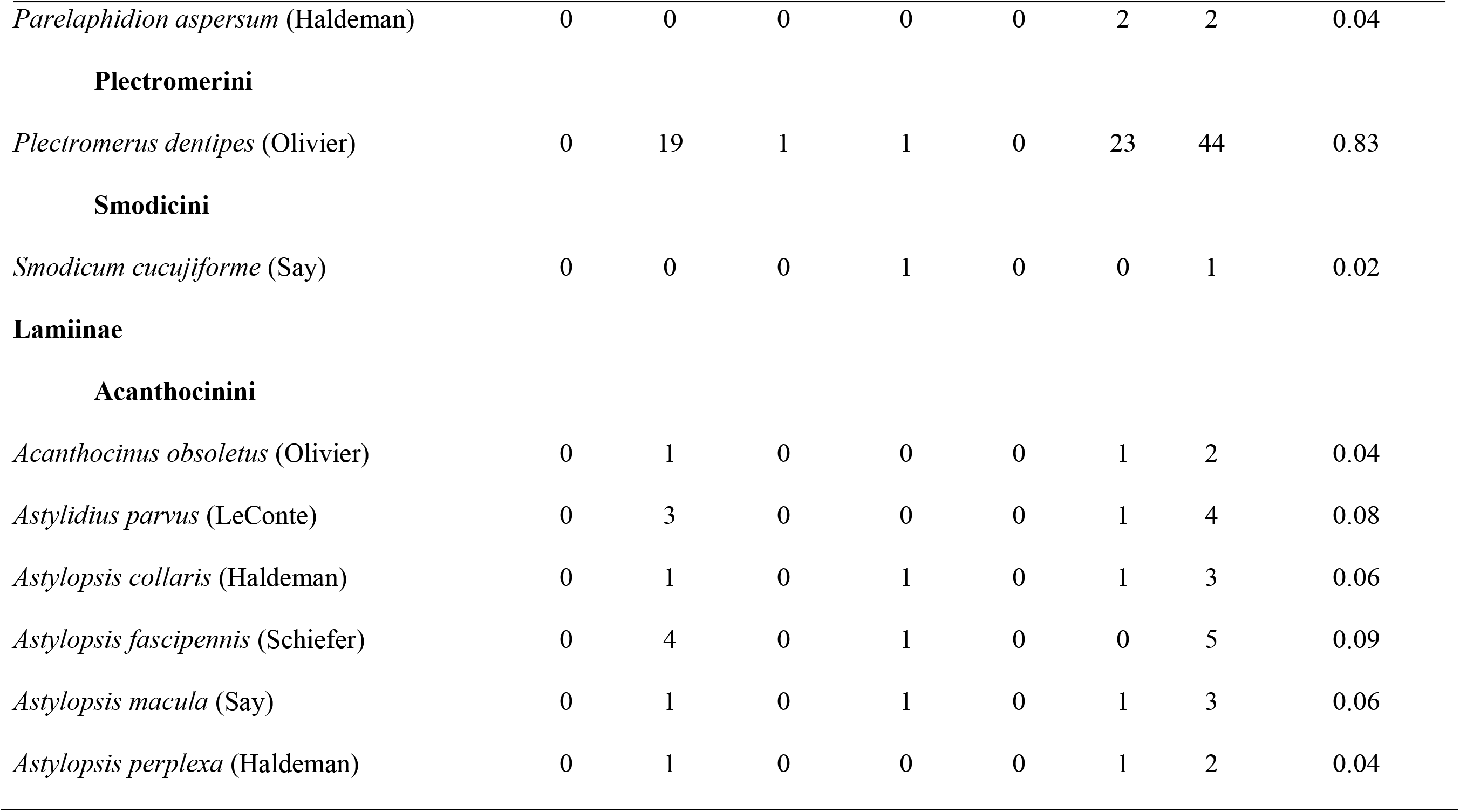
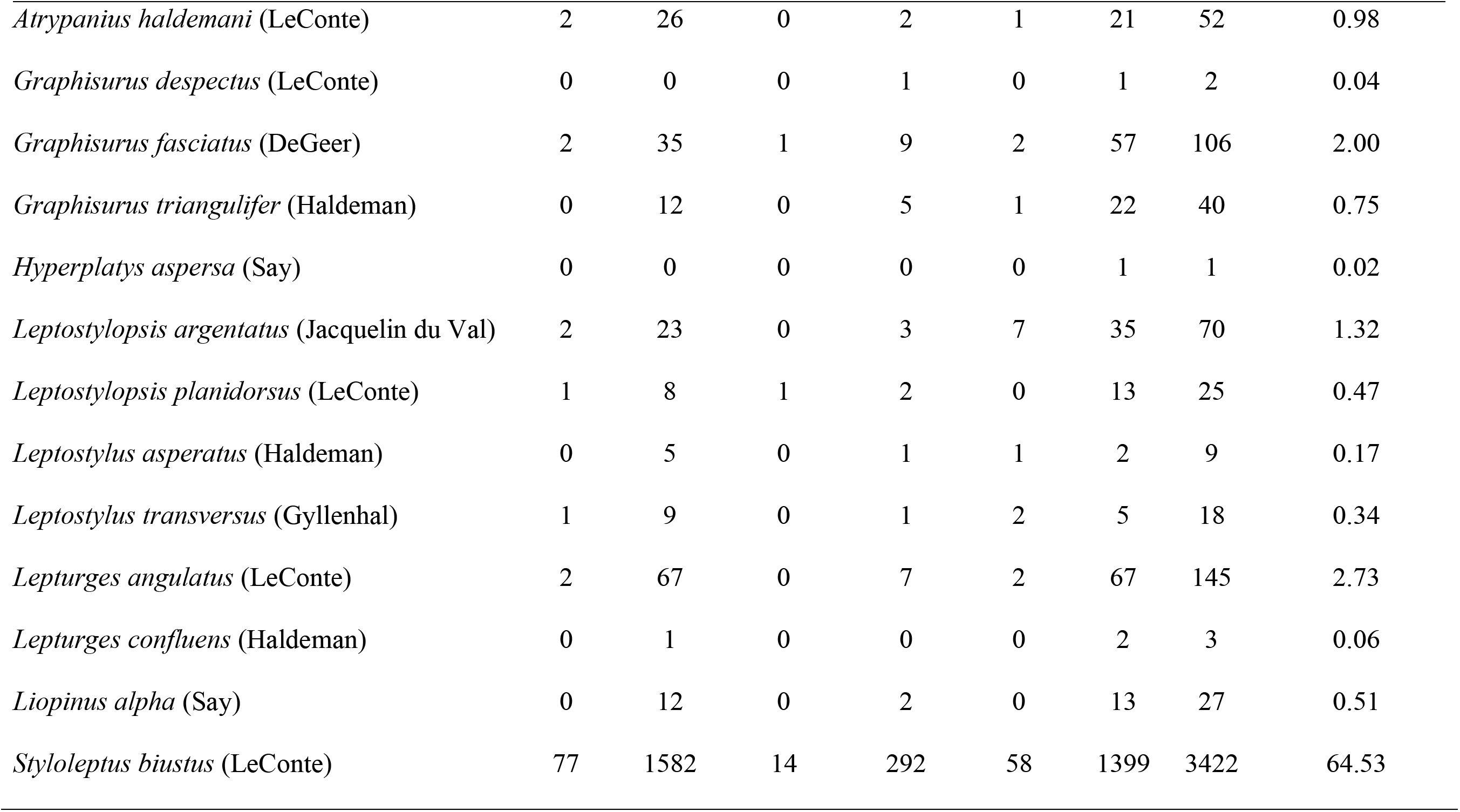
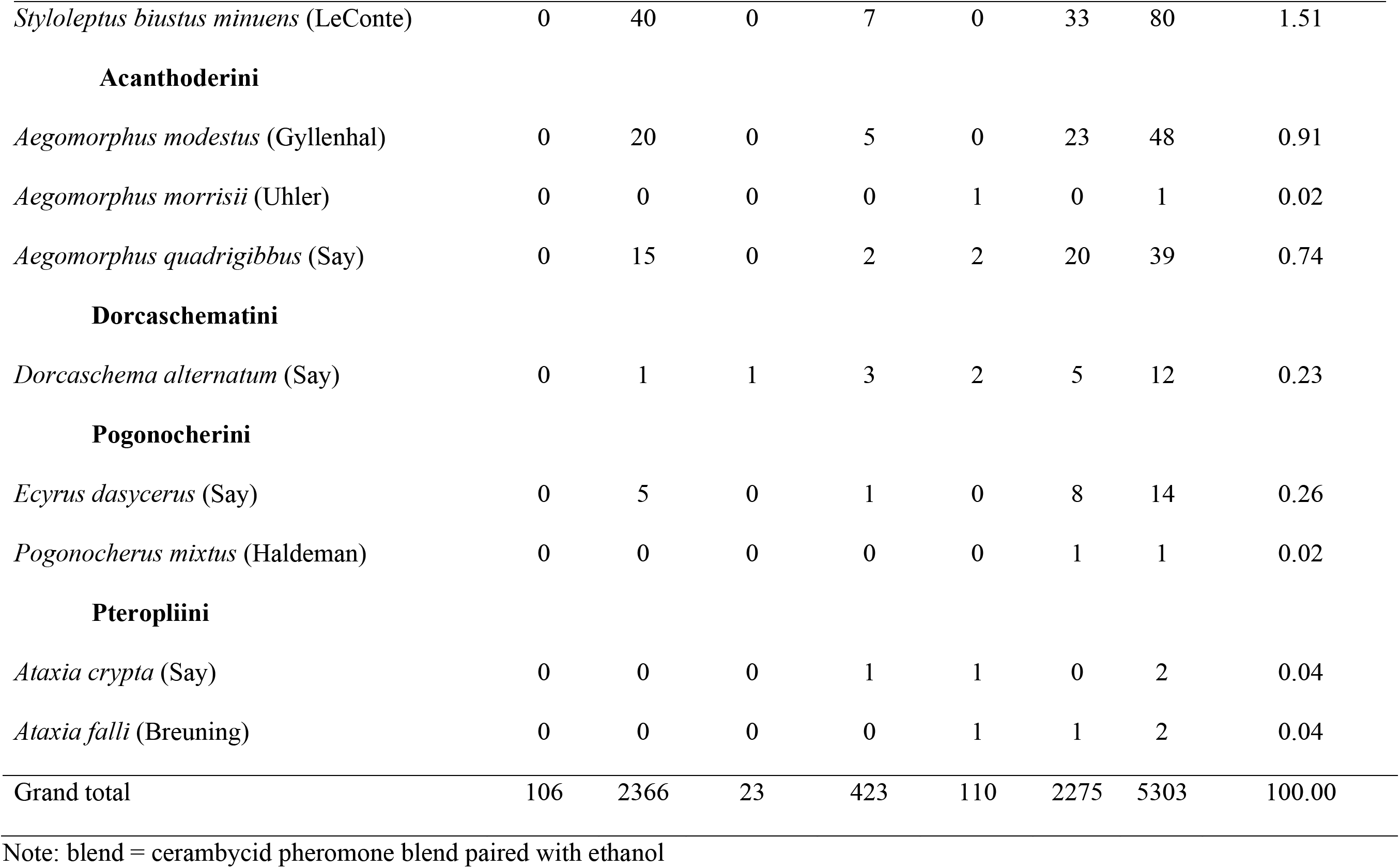
Taxonomy and number of cerambycid beetle species captured in Synergy multi-trap system (MTS) with and without Fluon application and Synergy funnel traps II (FT-II) baited with either low release ethanol lure or cerambycid pheromone blend paired with low release ethanol lure.

Forty-eight percent of the species we collected (21 out of 44) were recorded in limited numbers, not exceeding 5 individuals, with a single specimen recorded for 7 species of cerambycids. The most abundant species was *Styloleptus biustus* LeConte (n = 3,422) representing approximately 65% of all beetles captured (Table 3). The remaining 35% included *Neoclytus mucronatus* Fabricius (n = 465), *Curius dentatus* Newman (n = 197), *Elaphidion mucronatum* Say (n = 193), *Xylotrechus colonus* Fabricius (n = 158), *Lepturges angulatus* LeConte (n = 145), and *Graphisurus fasciatus* Degeer (n = 106) among the most frequently collected beetles (Table 3).

We captured 58.68% (n = 3,112) of longhorned beetles in the bottomland hardwood ecosystem (Audubon) and 41.32% (n = 2,191) from the loblolly pine dominant forest (Idlewild) (Supp. Table S3). At each site, 16 species of cerambycids were captured in ethanol baited traps, while 32 and 35 species of cerambycids were captured in traps with our cerambycid pheromone blend paired with ethanol baited traps, at Audubon and Idlewild respectively. Of the 44 species captured, 9 were captured exclusively in Idlewild and 8 in Audubon (Supp. Table S3); *Distenia undata* Fabricius, *Mallodon dasytomus* Say, *Orthosoma brunneum* Forster, *Enaphalodes atomarius* Drury, *Knulliana cincta cincta* Drury, *Neoclytus scutellaris* Olivier, *Smodicum cucujiforme* Say, *Astylopsis collaris* Haldeman, and *Leptostylus asperatus* Haldeman were only captured in pine dominated Idlewild, whereas *Elaphidion tectum* LeConte, *Parelaphidion aspersum* Haldeman, *Acanthocinus obsoletus* Olivier, *Astylidius parvus* LeConte, *Astylopsis perplexa* Haldeman, *Ataxia crypta* Say, *Ataxia falli* Breuning, and *Dorcaschema alternatum* Say were collected only in the bottomland hardwood swamp of Audubon (Supp. Table S3). Statistically higher numbers of *Curius dentatus* Newman, *Neoclytus mucronatus* Fabricius, *Plectromerus dentipes* Olivier, *Aegomorphus quadrigibbus* Say, *Ecyrus dasycerus* Say, *Leptostylopsis planidorsus* LeConte, *Lepturges angulatus* LeConte, *Atrypanius haldemani* LeConte, *Styloleptus biustus* LeConte, and *Styloleptus b. minuens* LeConte were collected in Audubon, whereas *Neoclytus scutellaris* Olivier, *Xylotrechus colonus* Fabricius, *Aegomorphus modestus* Gyllenhal, and *Graphisurus fasciatus* DeGeer were captured in statistically higher numbers in Idlewild (Supp. Table S4).

### Impacts of lure composition on capturing cerambycids

We supported our hypothesis that lure composition would impact the capture of longhorned beetles in our study. Seventeen species of longhorned beetles were significantly attracted to our blend of cerambycid pheromones paired with ethanol (Table 2). When Synergy MTS and FT-II were baited with an ethanol lure, they captured 2.58 ± 0.47 (mean ± SE) and 2.67 ± 0.33 cerambycids per trap per date respectively (Fig. 3). But, when our traps were baited with a blend of cerambycid pheromones plus ethanol, the mean number of cerambycids captured per trap per date increased drastically by 10.7X, with MTS and FT-II capturing 20.88 ± 2.12 and 32.48 ± 3.12 number of beetles per trap per date respectively (β = 2.37 ± 0.10 SE, z = 24.78, p < 2*10^−16^; Fig. 3).

**Figure 3.**
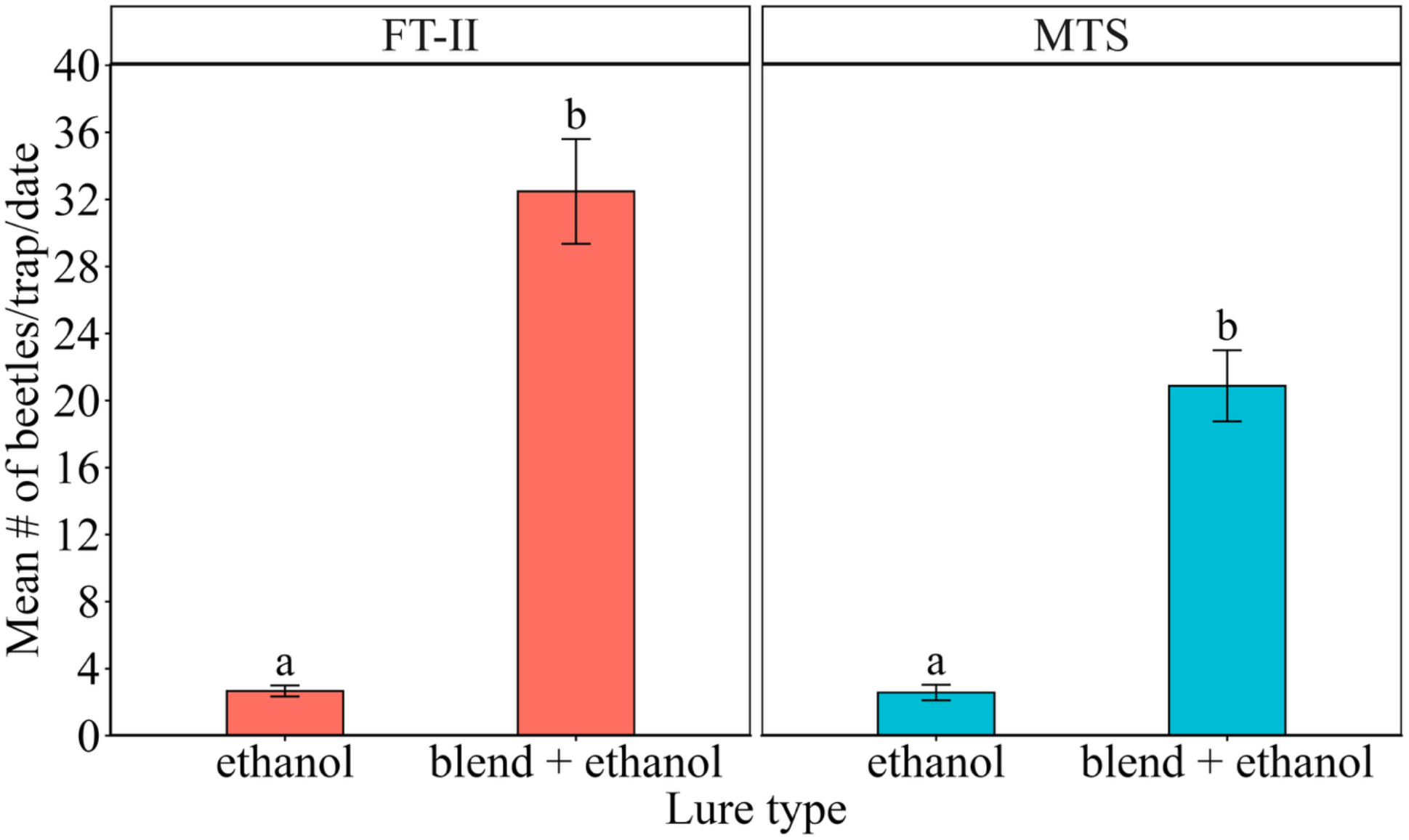
Mean (± SE) number of cerambycid beetles captured per trap per date in Fluon treated Synergy Funnel Trap II (FT-II) and Synergy Multi-trap System (MTS) with and without Fluon application. Traps were baited with either low release ethanol lure or blend of cerambycid pheromones paired with low-release ethanol lure. Different lowercase letters indicate significant differences among lure types at *P* < 0.01 level (Tukey’s HSD test).

### Impact of trap type on capturing cerambycids

We found partial support for our hypothesis that trap type would influence the capture of longhorned beetles in our study. Only 3 species of beetles were significantly affected by trap type, with higher capture of *Curius dentatus* Newman and lower capture of *Graphisurus fasciatus* DeGeer and *Leptostylopsis argentatus* du Val in MTS compared to FT-II (Table 2). However, when captures were aggregated across all species, the overall capture efficiency did not differ between trap types (Fig. 4). When comparing the Fluon treated Synergy MTS and FT-II, both were found to be equally effective, capturing 23.07 ± 2.66 and 21.71 ± 2.43 number of longhorned beetles per trap per date respectively (β = 0.01 ± 0.07 SE, z = 0.12, p = 0.904545; Fig. 4).

**Figure 4.**
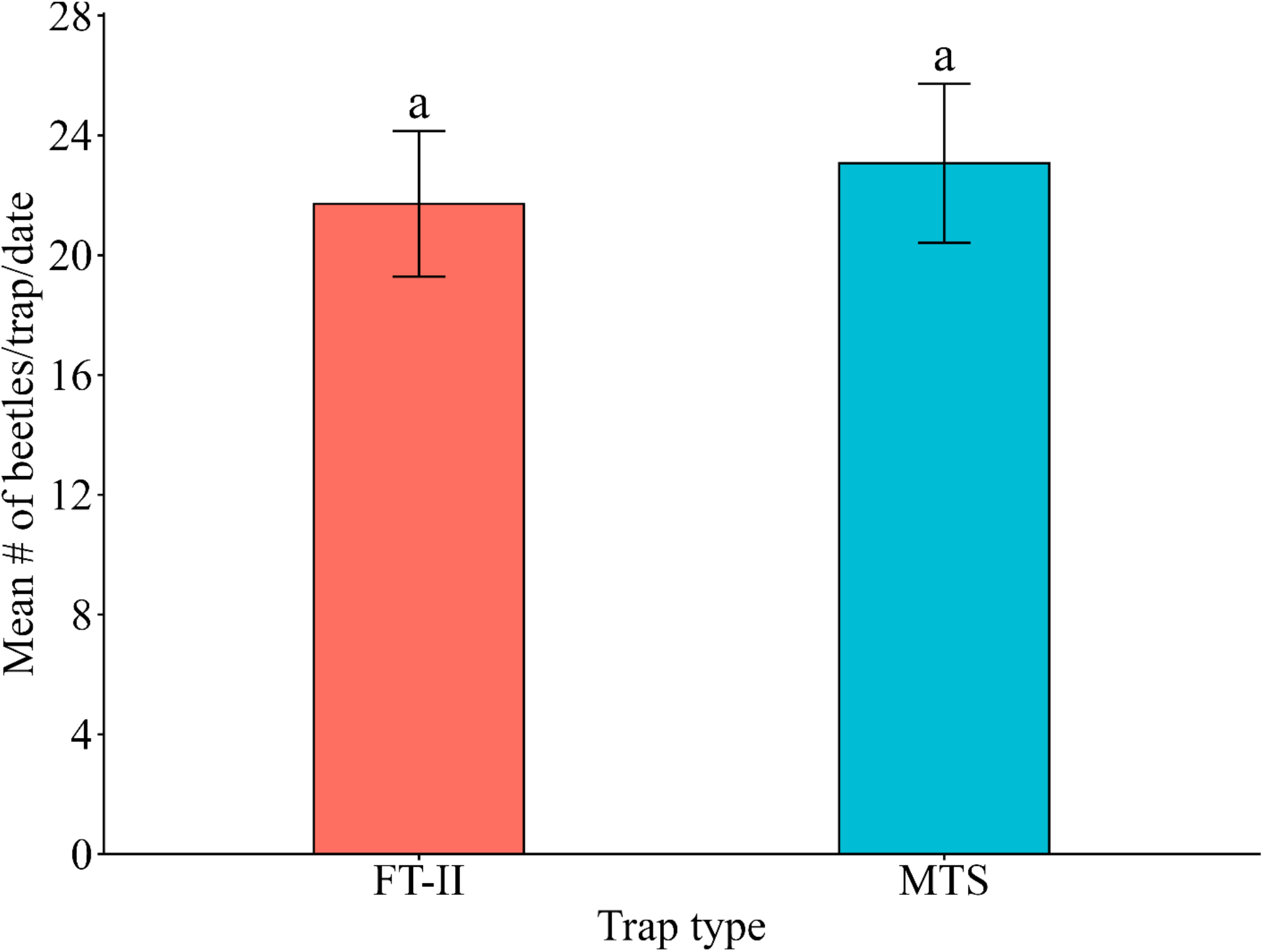
Mean (± SE) number of cerambycid beetles captured per trap per date in Synergy Funnel Trap II (FT-II) and Synergy Multi-trap System (MTS) both treated with Fluon. Traps were baited with either low release ethanol lure or blend of cerambycid pheromones paired with low-release ethanol lure. Same lowercase letters indicate no significant differences among trap types.

### Impact of Fluon application on capturing cerambycids

We failed to support our hypothesis that Fluon application wouldn’t impact the capture of longhorned beetles in the Synergy MTS. The application of Fluon to Synergy MTS significantly increased the captures of 15 species of beetles compared to untreated Synergy MTS (Table 2). Fluon treated Synergy MTS captured 23.64 ± 2.57 longhorned beetles per trap per date, which is 4.30X greater than MTS without Fluon, which captured 5.79 ± 0.80 beetles per trap per date (β = 1.46 ± 0.11 SE, z = 13.54, p < 2*10^−16^; Fig. 5).

**Figure 5.**
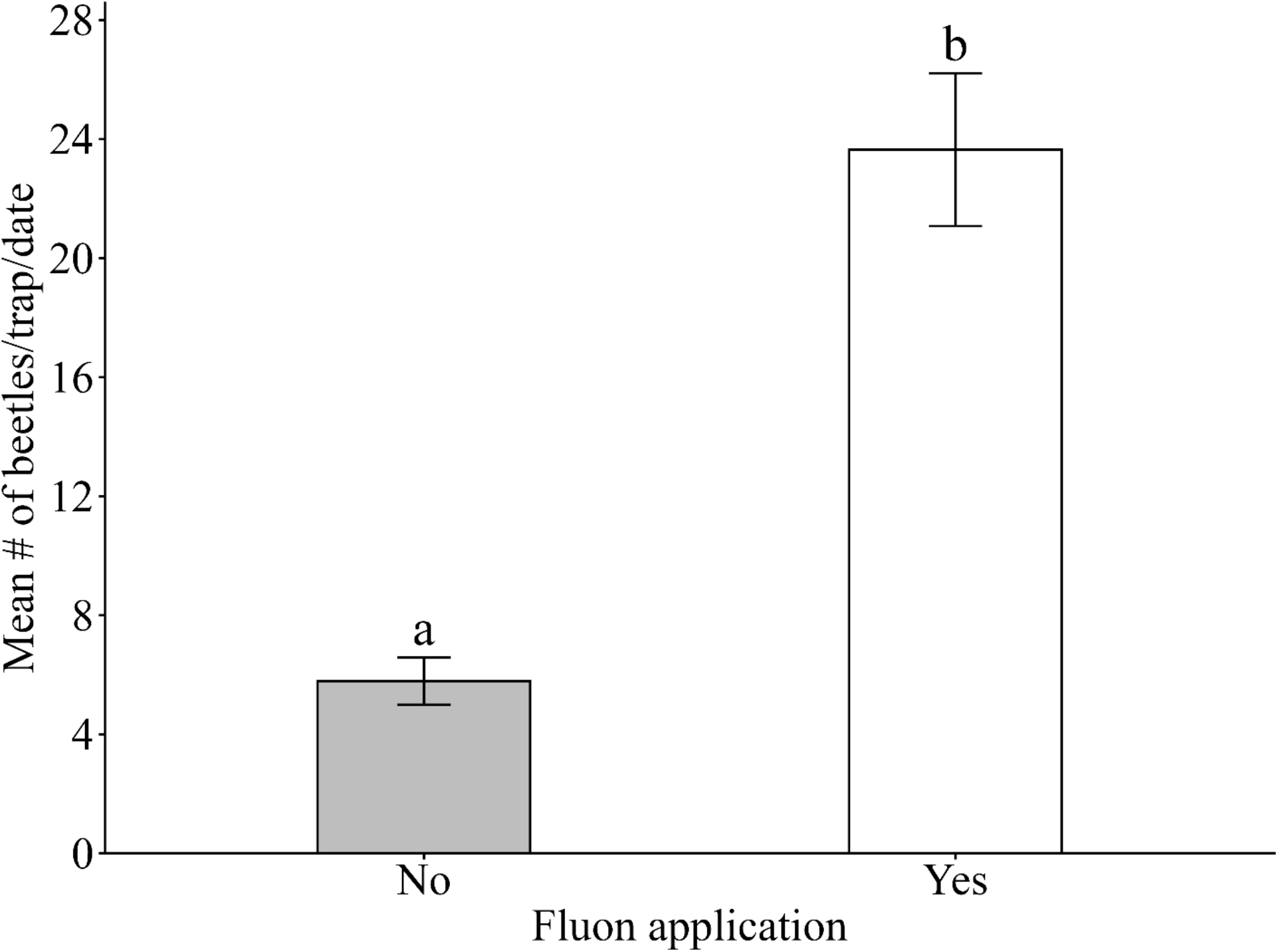
Mean (± SE) number of cerambycid beetles captured per trap per date in Synergy Multi-trap System (MTS) with and without Fluon application. Traps were baited with either low release ethanol lure or blend of cerambycid pheromones paired with low-release ethanol lure. Different lowercase letters indicate significant differences among Fluon treated vs untreated MTS at *P* < 0.01 level (Tukey’s HSD test).

## Discussion

### Chemical composition of lures played a critical role in capturing target and non-target longhorned beetles

While the inspiration for our research was to evaluate if the Synergy MTS traps required an application of Fluon to effectively capture target species of longhorned beetles (i.e., those known to be attracted to pheromones), the inclusion of an ethanol lure on our control traps allowed us to also evaluate the impact of lure on the response of longhorned beetles. We found that by adding a blend of cerambycid pheromones to a trap with an ethanol lure, total captures of longhorned beetles increased by 10.7X, as compared to traps baited with an ethanol lure alone, supporting our hypothesis that lure composition would influence the capture of longhorned beetles (Fig. 3). In total, 17 species of longhorned beetles were captured in significantly higher numbers in traps baited with a blend of cerambycid pheromones paired with an ethanol lure, as compared to an ethanol lure alone (Table 2). This is not a surprising result, as components within the blend of cerambycid pheromones deployed in our study are known, singly or in combination, to attract a broad diversity of cerambycid beetles within subfamilies Cerambycinae, Lamiinae, and Spondylidinae (Halloran et al., 2018; Hanks et al., 2018; Hanks et al., 2012).

### Validation of known attractants and opportunities for broadening and deepening our understanding of the chemical ecology of longhorned beetles

Among the 44 species of longhorned beetles captured, 18 species have previously documented aggregation-sex or sex pheromones (i.e., volatile chemicals produced by beetles with field bioassays demonstrating their attractive function), 11 species have known attractants (i.e., volatile chemical compounds used in field traps that elicit orientation or attraction responses in the beetles, but have not yet been confirmed as naturally produced pheromone through laboratory analyses such as headspace volatile collections), and 15 species have no information on their semiochemicals (i.e., no previously documented attractants or pheromones; Table 4). Among the 17 species significantly attracted to our blend of cerambycid pheromones paired with an ethanol lure, we validated the attractive responses of 11 species to one or more of the components of our pheromone blend—*Acanthocinus obsoletus*, *Aegomorphus quadrigibbus*, *Astylopsis collaris*, *Ataxia crypta*, *Curius dentatus*, *Dorcaschema alternatum*, *Eburia quadrigeminata*, *Ecyrus dasycerus*, *Graphisurus despectus*, *Leptostylus asperatus*, and *Styloleptus biustus biustus*—in a novel geographic and climatic context. These species had not previously been evaluated in subtropical southeastern Louisiana, where our findings confirm that their pheromone-mediated attraction extends to this region’s distinct forest ecosystems and hot and humid conditions.

**Table 4.**
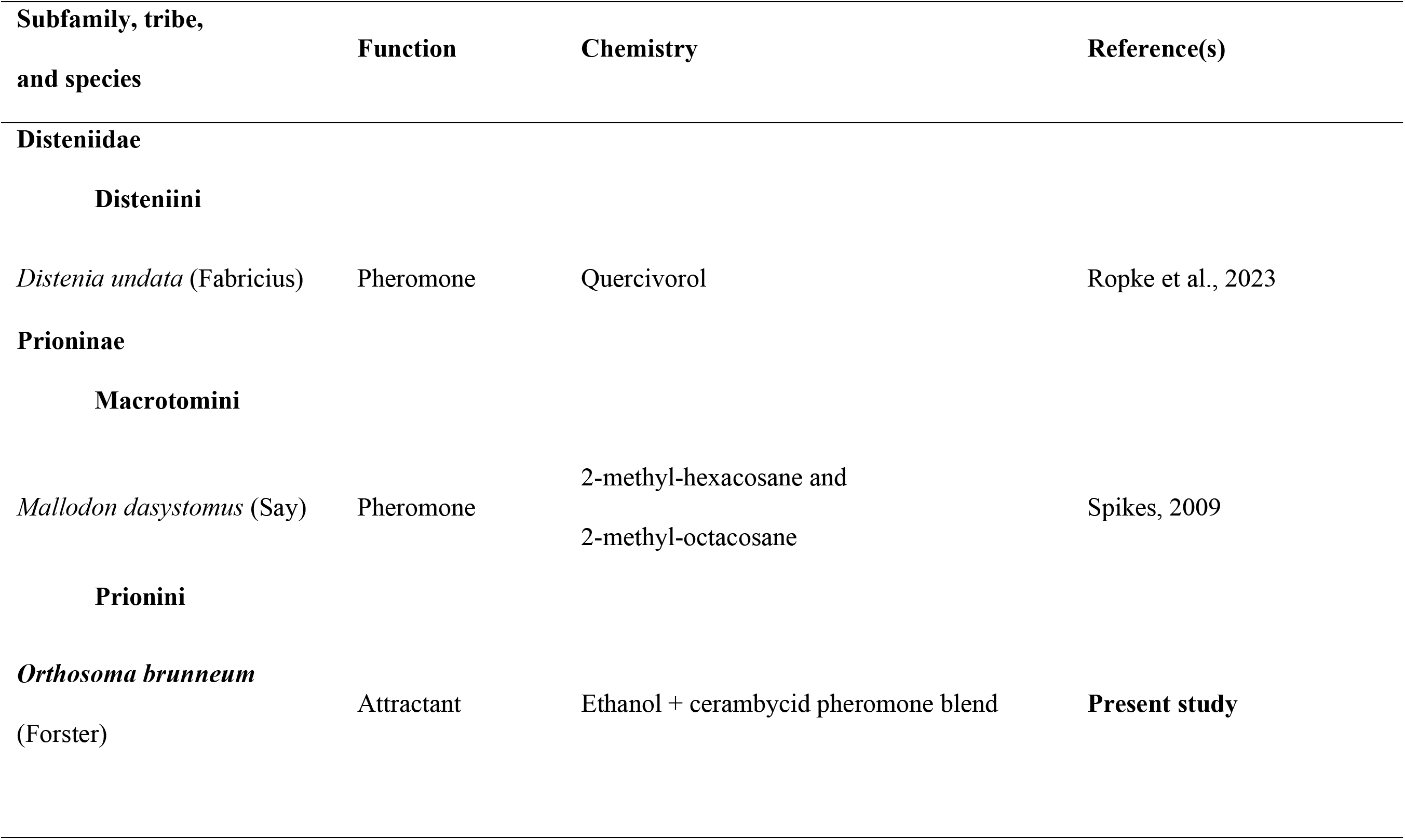
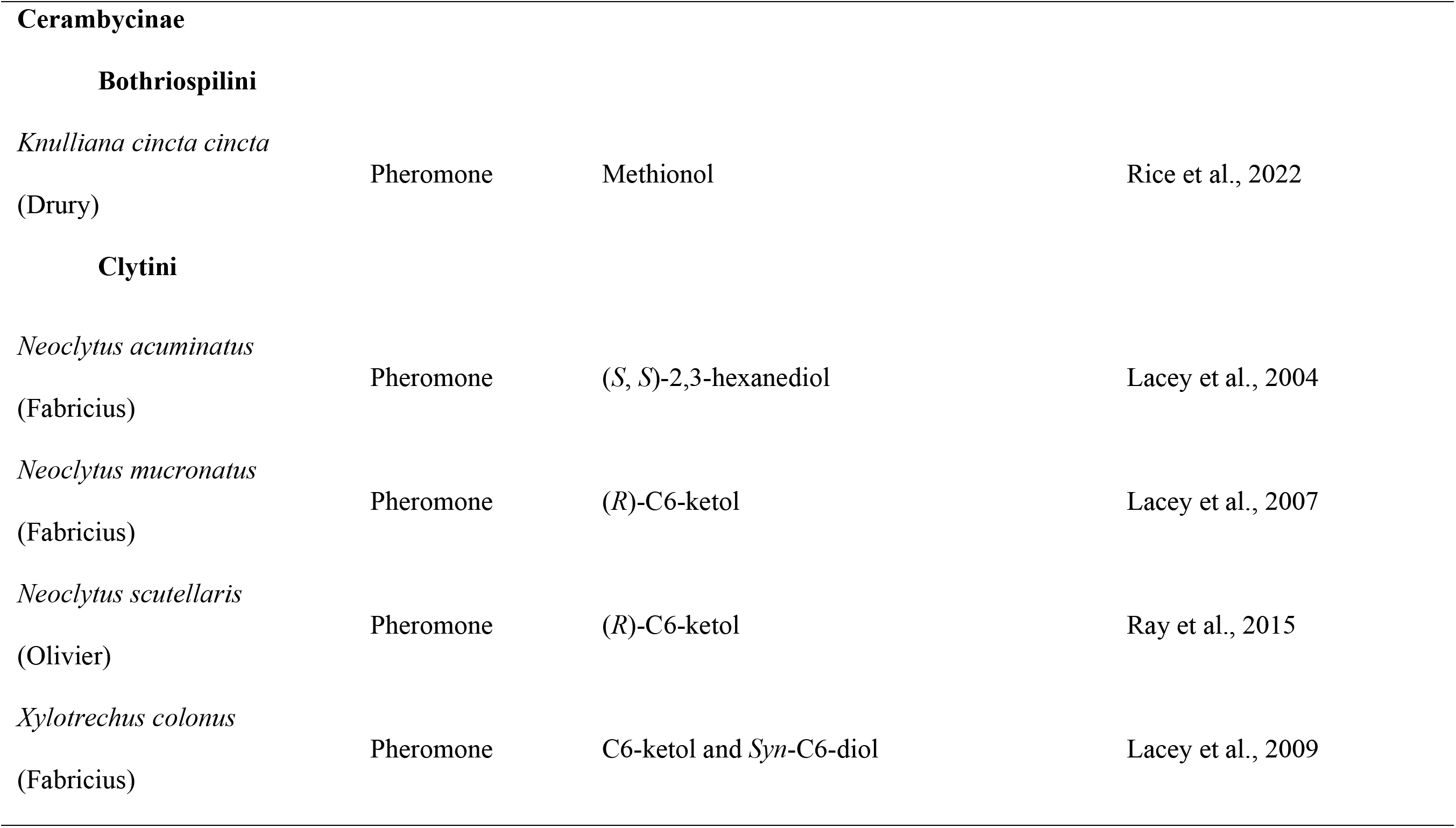
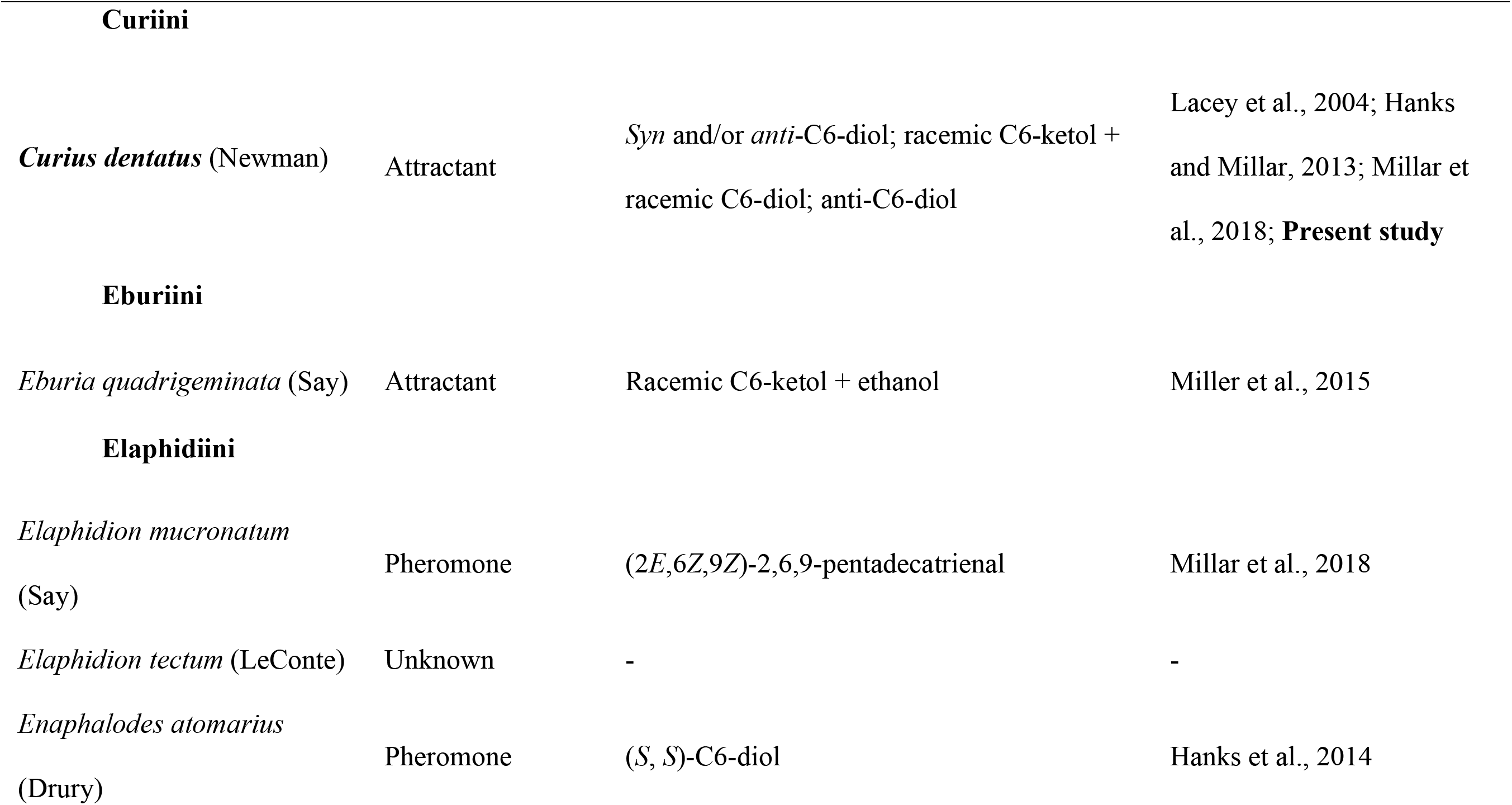
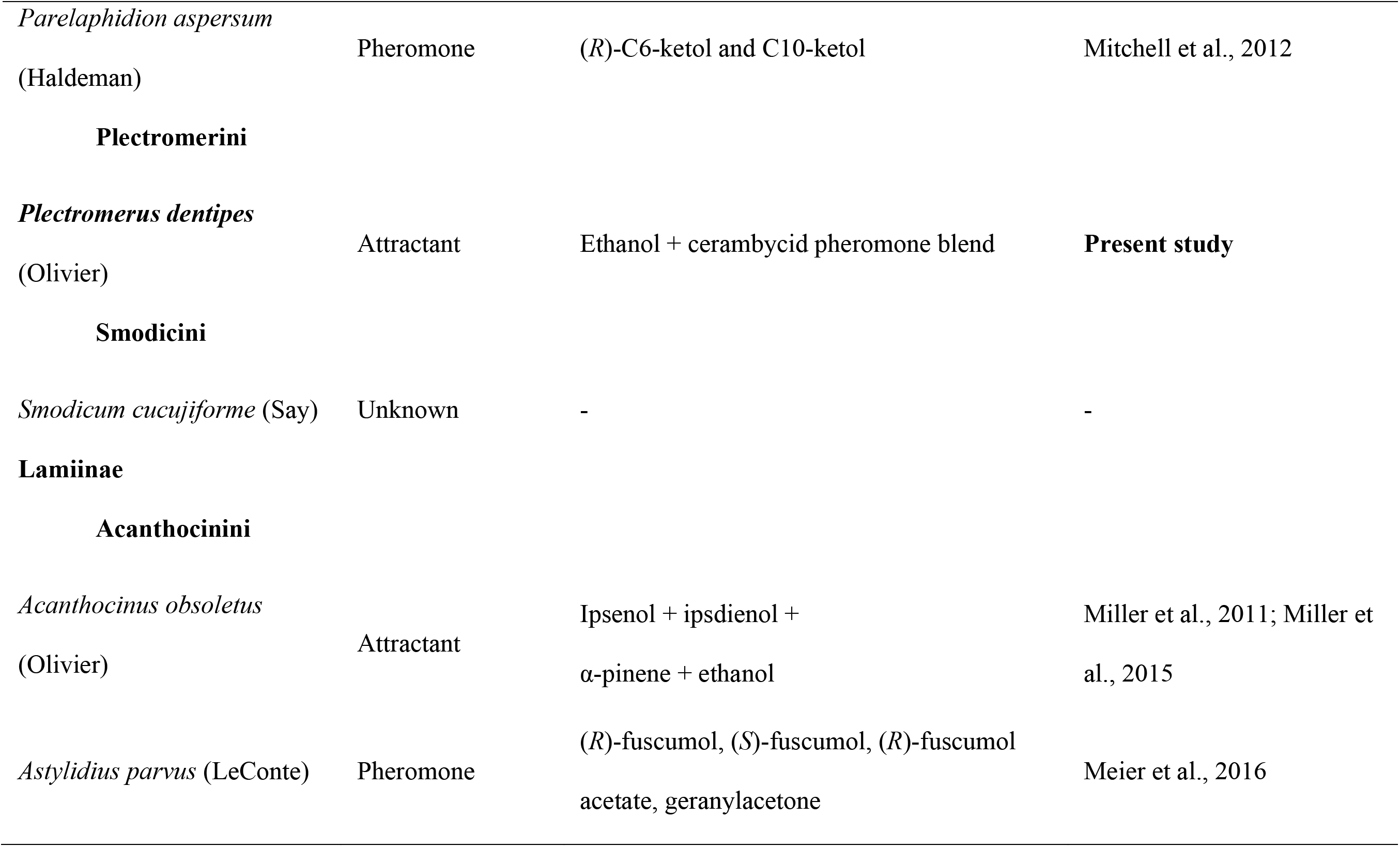
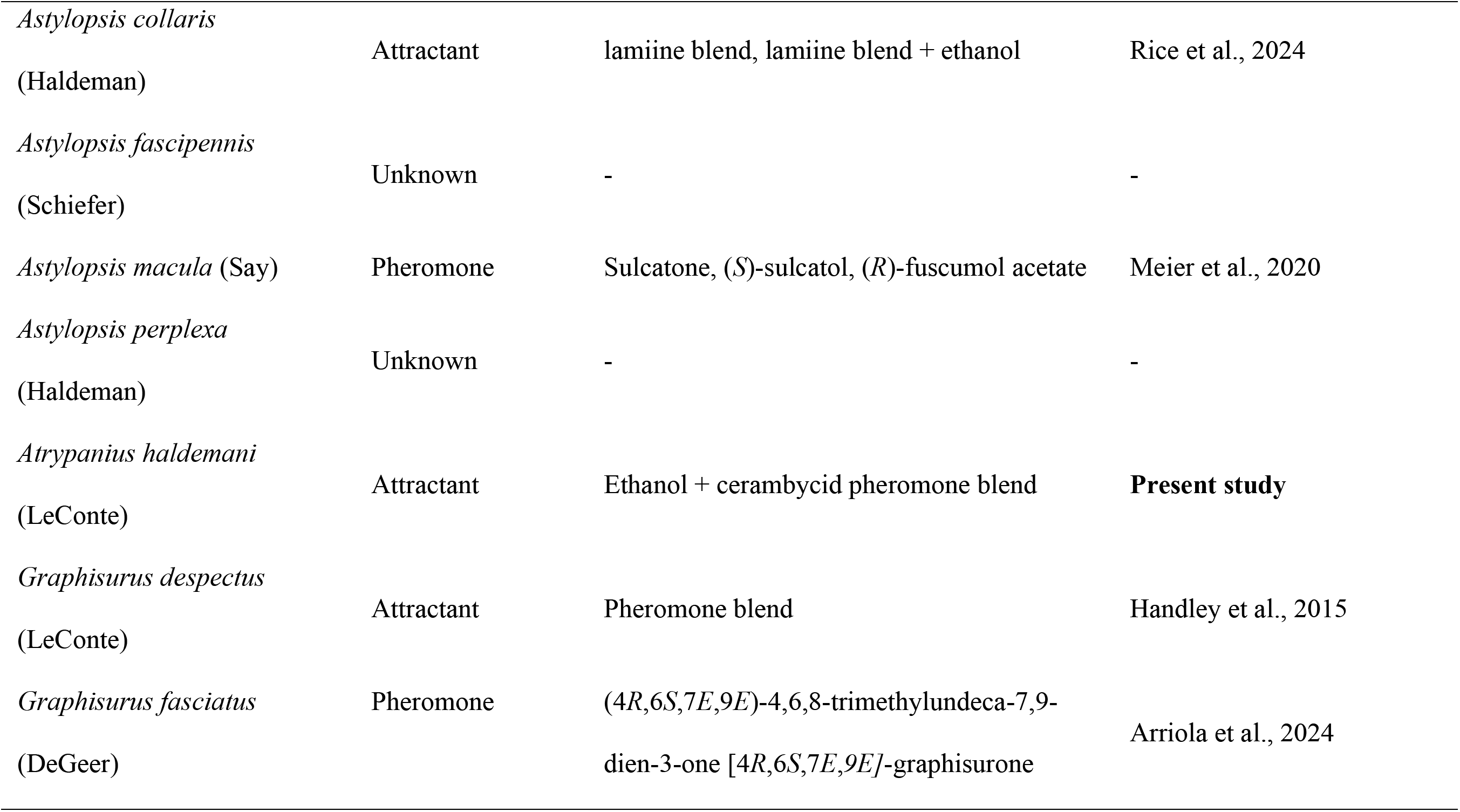
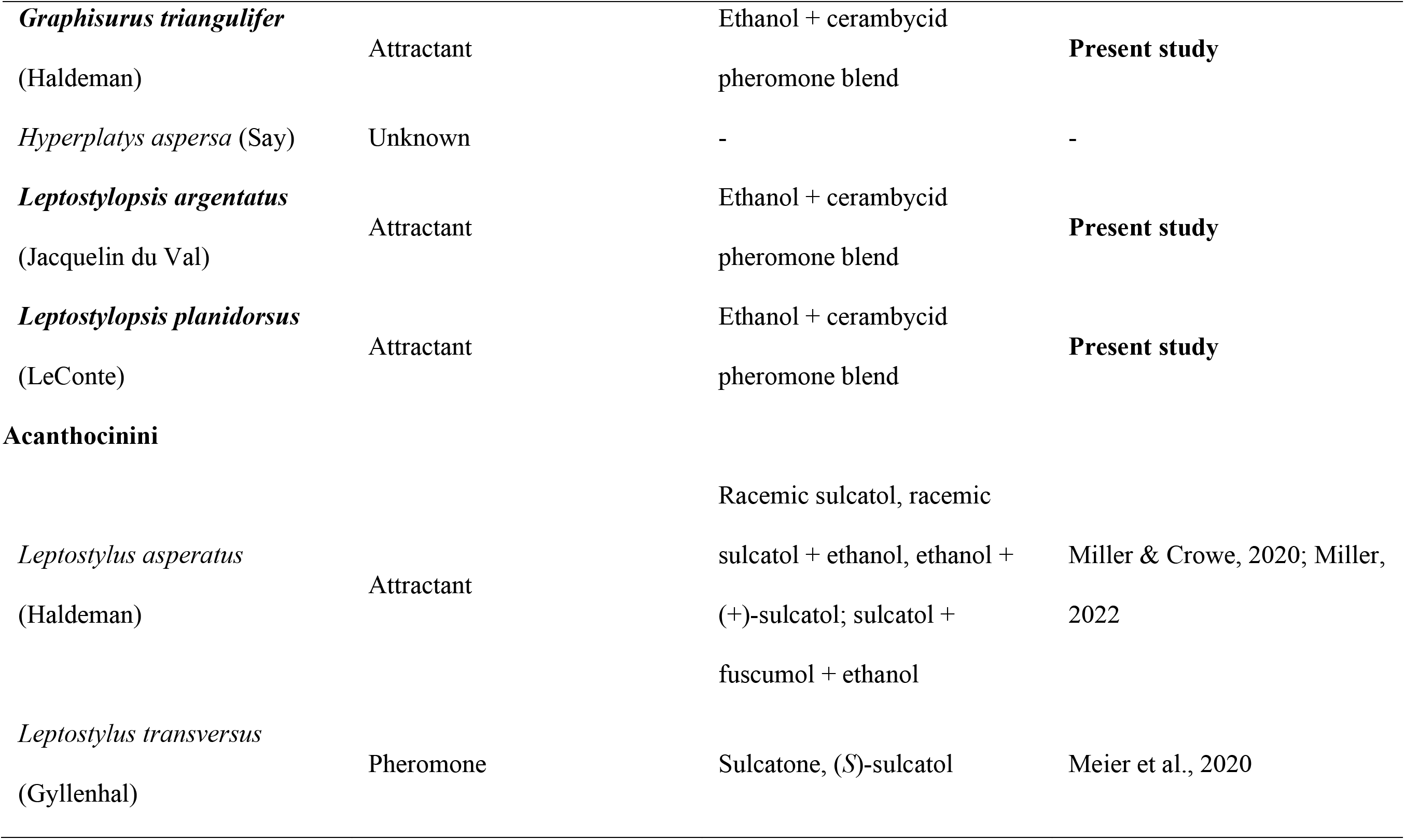
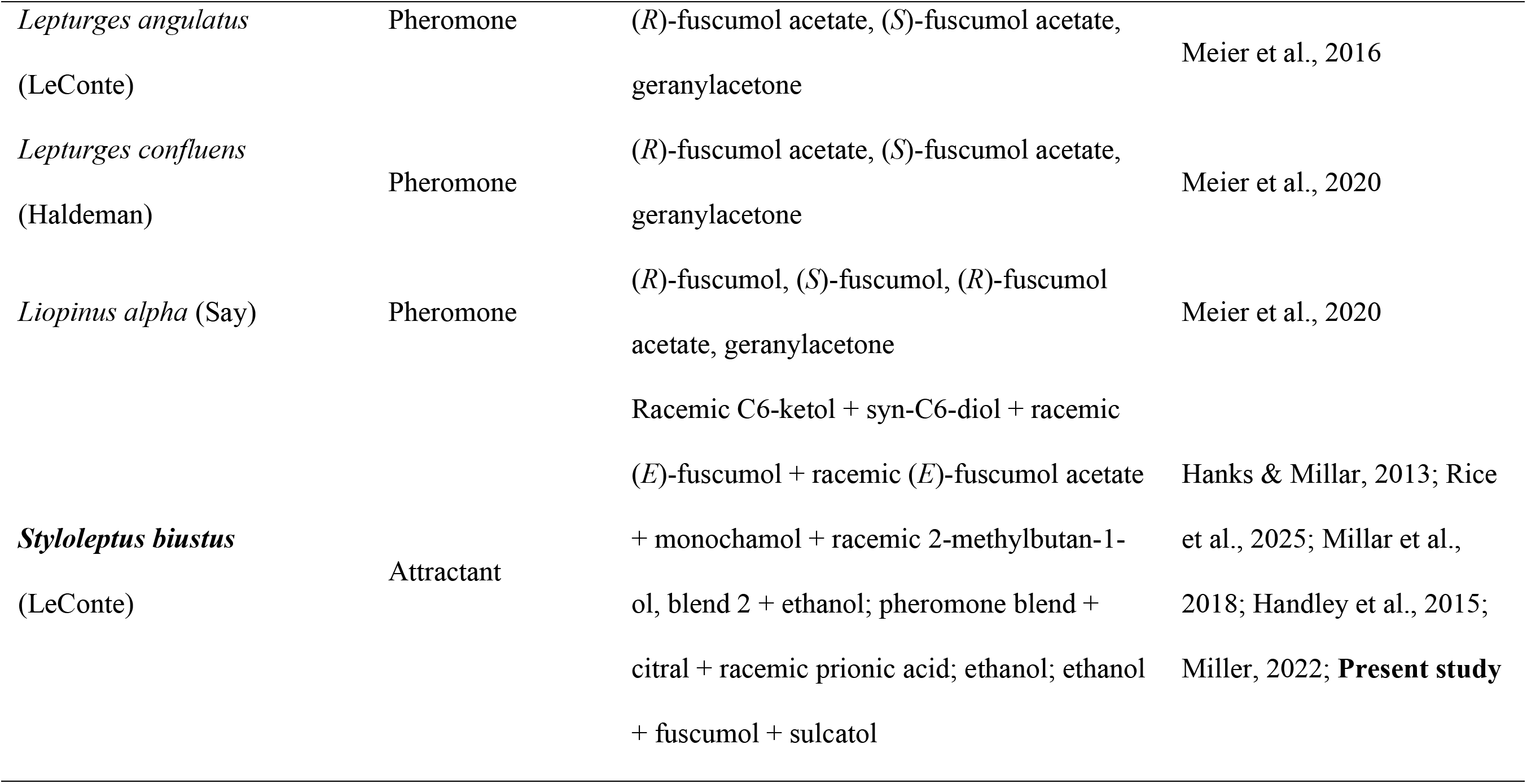
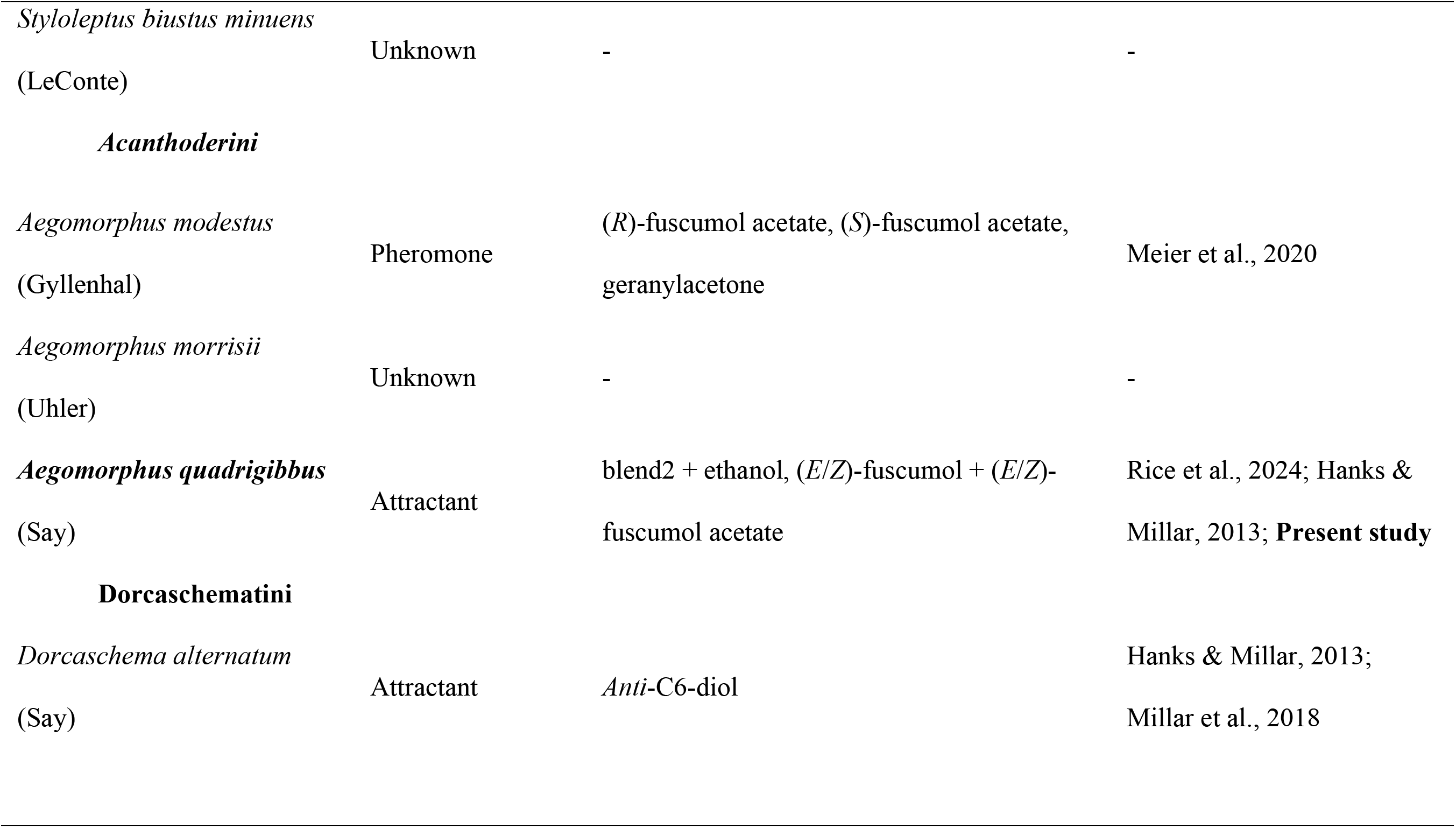
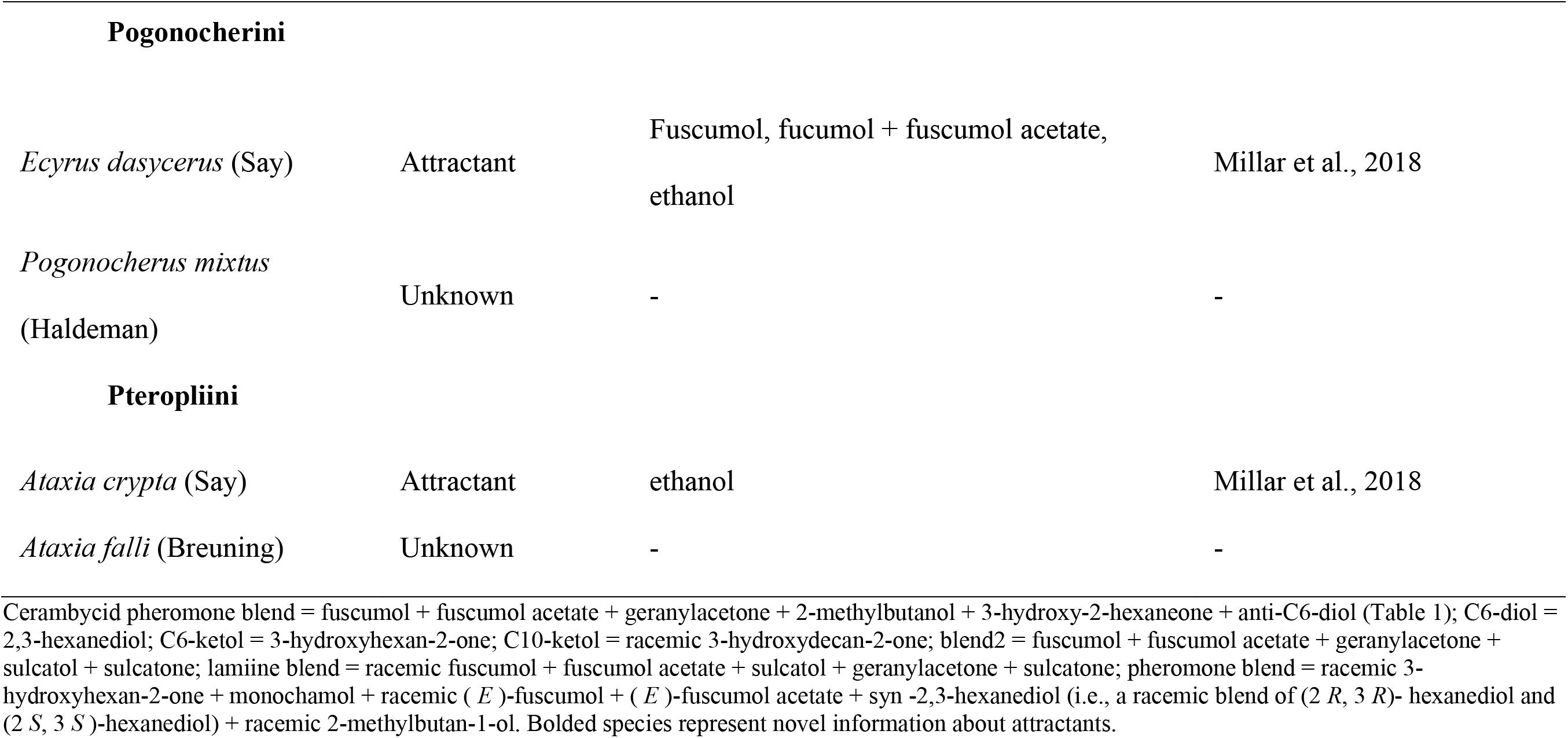
Chemistry of pheromones and attractants for cerambycid species that were captured at two distinct sites: Audubon and Idlewild in southeastern Louisiana.

Notably, we added information on attractants for 6 species of longhorned beetles, Prioninae: *Orthosoma brunneum*, Cerambycinae: *Plectromerus dentipes*, and Lamiinae: *Leptostylopsis argentatus*, *Atrypanius haldemani*, *Leptostylopsis planidorsus*, and *Graphisurus triangulifer*, all of which were found to respond significantly to our cerambycid pheromone blend paired with ethanol (Table 2). Three of these species, *L. argentatus*, *L. planidorsus*, and *G. triangulifer*, occur in genera that have previously described attractants and pheromones. This pattern is consistent with the high degree of semiochemical parsimony observed within longhorned beetles, wherein closely related species may share major pheromone components, or even produce identical compositions of pheromones, but still avoid cross-attraction. Such specificity is maintained through pre-zygotic reproductive isolation mechanisms that operate spatially (i.e, differences in geographic range or microhabitat use), temporally (i.e., seasonal phenology or non-overlapping diel cycles of pheromone production and activity), and behaviorally (i.e., species-specific responses to pheromone or host volatiles), thereby preventing interspecific mating interference (Hanks & Millar, 2013; Mitchell et al., 2015). For example, other species of longhorned beetles in the genus *Leptostylus* have been previously shown to respond to traps baited with the pheromone fuscumol, a common component of blends of pheromones produced by members of the Lamiinae tribes Acanthocinini and Acanthoderini (Hanks & Millar, 2013; Mitchell et al., 2011). There is currently no evidence to support fuscumol by itself as a pheromone for beetles in the genus *Leptostylus*. Regardless, pheromones of different species of longhorned beetles may act as kairomones that aid in locating reproductive and oviposition opportunities (Collignon et al., 2016; Reddy & Guerrero, 2004; Yang et al., 2004). Future research should evaluate the role of fuscumol as an attractant for beetles in the genus *Leptostylus*.

Herein, we also provided evidence for our cerambycid pheromone blend as the first reported attractants for *Plectromerus dentipes* (Cerambycinae: Plectromerini) and *Orthosoma brunneum* (Prioninae: Prionini). Similar to the acanthocinines, these findings may represent additional cases of semiochemical parsimony within the Cerambycidae. Previously described as a member of the Curinii (Nearns & Branham, 2008), it is possible that *P. dentipes* shares an attraction to the same compound as *Curius dentatus*, *anti*-2,3-hexanediol, which was included as part of our cerambycid pheromone blend. Similarly, *O. brunneum* may also be attracted to *anti*-2,3-hexanediol, as reported for *Tragosoma pilosicorne* Casey (Ray et al., 2012). Lastly, we found that *S. biustus* responded significantly to our cerambycid pheromone blend while *S. b. minuens* did not (Table 2). Such apparent difference in response may be the result of substantially lower number of *S. b. minuens* captured (n = 80) compared to *S. biustus* (n = 3422) potentially reflecting limited statistical power rather than a true biological difference. Thus, additional sampling would be required to determine if these closely related taxa differ in their pheromone ecology at the subspecies level. Altogether, these finding highlight the need for future subtractive field studies (i.e., testing all combinations of pheromone blends together and separate) combined with the collection of pheromones from live adult male beetles to identify and validate the pheromones produced by these collected beetles (Hanks et al., 2019).

### The Synergy Multi-Trap System and Synergy Funnel Trap II were equally effective in capturing longhorned beetles

Contrary to our hypothesis that trap type would influence the collection of cerambycids, we found that Fluon treated Synergy MTS and FT-II were equally effective in capturing longhorned beetles (Fig. 4). The comparable efficiency of both traps may be explained by their shared design features including black color, vertical orientation, wet collection cups, and nearly equal interception surface area (MTS = 0.72 m^2^ vs FT-II = 0.84 m^2^) that beetles encounter during flight. Despite using different formulations of plastics when manufacturing the Synergy MTS and FT-II (Personal Communication with R. Setter), application of Fluon may have altered the surface textures of both traps, leading to equivalent rates of beetle capture. Our result aligns with the findings of Allison et al. (2014), who also reported no difference in the capture of Cerambycinae and Lamiinae beetles between Teflon or Fluon-treated multiple-funnel vs panel traps, or untreated multi-funnel vs panel traps that were produced by a different manufacturer.

### Materials used to manufacture Synergy Multi-Trap System did not eliminate the need for Fluon application to enhance longhorned beetle collection

When Synergy MTS without an application of Fluon were deployed in the field, they captured relatively few beetles. However, capture rates of longhorned beetles increased by 4.3X when Fluon was applied to MTS under identical conditions (Fig. 5). This finding contrasts with our hypothesis that the material used to manufacture MTS would eliminate the need for Fluon application. Our results suggest that the material lacked the chemical or structural properties necessary to create a sufficiently hydrophobic, slippery, or low friction surface, or that these properties were diminished during the manufacturing process through mixing with other compounds (Peisker et al., 2014). Low material viscosity or rapid evaporation may have also reduced surface slipperiness over time (Gorb, 2001; Peisker & Gorb, 2012). As a result, beetles may have been able to alight on, cling to, or crawl across the traps and ultimately escape (Geiselhardt et al., 2009; Geiselhardt et al., 2010).

Application of Fluon on MTS may have further increased surface slipperiness, resulting in higher beetle captures. This is because Fluon is thought to decrease the ability of intercepted beetles to alight on, cling to, or crawl across Fluon-treated trap surfaces, thereby increasing the likelihood that they fall into attached collection cups. Our findings align with previous studies demonstrating enhanced captures of longhorned beetles in Fluon treated funnel traps equipped with wet collection cups, as well as Fluon applied panel traps with dry or wet collection cups (Graham et al., 2010; Graham & Poland, 2012). While largely understood to function as a friction reducer, how exactly Fluon application impacts interactions between longhorned beetle tarsi and trap surfaces remain poorly understood. Furthermore, it is unclear whether the effectiveness of Fluon varies between species of longhorned beetles according to their natural history (e.g., feeding guild, oviposition preferences) and warrants further investigation.

### The structure of forest ecosystems may influence the capture of unique and abundant longhorned beetles

During our study, we captured 3,112 beetle specimens (58.68%) at the Audubon Nature Center— a 35-hectare bottomland hardwood forest, compared to 2,191 specimens (41.32%) at the 526-hectare loblolly pine dominated Idlewild Research Station (Supp. Table 3). At Audubon, 32 species were captured with our blend of cerambycid pheromones paired with ethanol vs 16 in only ethanol baited traps, whereas at Idlewild the number was 35 vs 16 respectively. Twelve species of cerambycids were captured only at Idlewild and eight at Audubon. These disparities in collection may stem from contrasting forest compositions, differences in total forested areas of our sampling sites or the structure of surrounding habitat. For example, the specialist *Orthosoma brunneum*, which relies on conifers (e.g., *Pinus*), was captured only at Idlewild—a finding consistent with site-specific lure impacts at this location (e.g., Lingafelter, 2007; Wong, 2016). However, the role of site in explaining the collection of non-target beetles (i.e., those not responsive to pheromone blends) is not clear. The most likely explanation is that beetles were randomly intercepted.

Our findings primarily describe the pheromone ecology of species of cerambycids targeted by our blend, rather than the entire fauna. Thus, non-detections of cerambycids not responsive to our pheromone blend should not be interpreted as absence at a site. Conversely, the low captures of *E. atomarius*, *A. collaris*, *L. asperatus*, and *G. despectus* at Idlewild and *A. crypta*, *A. obsoletus*, *A. parvus*, *D. alternatum*, and *P. aspersum* at Audubon whose pheromones fall under the taxonomic coverage of our pheromone blend, may indicate their local rarity or that our field bioassay was executed outside their flight period.

### Testing and evaluating hypotheses of trap efficacy with target as well as non-target species of woodboring beetles

The most effective strategy to reduce potential ecological and economic damage after the introduction of an invasive species is eradication of its population before they can grow and spread (Lodge et al., 2006; Myers et al., 2000). Because many species of woodboring insects are often difficult to detect and responsive to long-range volatile attractants, considerable efforts have been invested into the development of traps that attract beetles across large distances, facilitating their rapid detection (Hanks & Millar, 2016). As woodborers approach traps from long distances, they integrate information provided by volatile chemicals along with other sensory modalities such as vision, to determine whether to land or not while at shorter distances (Besana et al., 2025; Cavaletto et al., 2021; Johnson et al., 2019). If woodborers choose to land, the three-dimensional structure and properties of the trap, including its trap surface, often influence the retention of target insects (Graham et al., 2010; Saint-Germain et al., 2007). Thus, researchers have sought to evaluate the contributions of multiple components in successfully capturing and retaining target species, as well as to develop multi-purpose traps that can be deployed to detect multiple target and non-target species simultaneously (Allison et al., 2014; Graham et al., 2010; Hanks & Millar, 2018; Hanks et al., 2012; Rice et al., 2020).

We argue that with respect to trapping studies, there are number of considerations, some of which are addressed and discussed herein with respect to the analysis of our own data, that can confound the ability of researchers to rigorously test and generate support for or against hypotheses regarding the response of woodboring insects to traps. The first consideration involves including non-target species in statistical evaluations of trap design, when researchers are using lures to explicitly or implicitly target specific species of woodborers for the purpose of assessing trap function. For instance, trap design had no significant influence on the capture of Cerambycidae when analysis was done at family level (p = 0.90; Fig. 4), but when analysis was conducted at the species level, the capture of three species were found to be impacted by trap design. Two factors are likely to drive these results; first, the most abundant target and non-target longhorned beetles are weighing the results of statistical analyses towards interpretations which may only apply to the specific circumstances and biology of those species. Of the nearly 37,000 species of cerambycids known, they exist across an enormous diversity of shapes, sizes, and behaviors. It is therefore not prudent to assume that responses by a few species of longhorned beetles reflect the potential responses by the remaining species.

Secondly, including non-target species into analyses with target species may introduce measurement error in an inconsistent fashion across the study unless specifically controlled for in the design (e.g., grids or multiple transects of unbaited traps). Further, in the case of target species that are rare or generally low in abundance, abundant non-target species may obscure an understanding of the factors driving their response, which may lead to Type I and Type II errors. Thus, while there certainly is an allure to making conclusions at the family level, we suggest that unless there is a compelling reason to do otherwise, each species of longhorned beetles be analyzed in separate statistical models, and then aggregated (i.e., Fig. 4), to improve assessments regarding the role of trap design on the collection of longhorned beetles for survey programs. Lastly, spacing between traps should also be taken into considerations when testing hypotheses of trap efficacy. Since the effective attraction distance of the lures used in this study is unknown, the 5 m spacing maintained between traps to maximize replication within the available study area may not have been sufficient to completely prevent interactions among neighboring traps. Overlapping odor plumes from adjacent or neighboring lures could have reduced treatment independence, potentially influencing beetle responses and obscuring true treatment effects. Thus, to ensure independent assessments of trap efficacy, future studies should evaluate the role of inter-trap distances on each of the pheromones deployed in our study to determine the optimal spacing required to minimize potential treatment interference.

### Fundamental and applied implications and directions for future studies

Our study demonstrated that traps baited with blend of cerambycid pheromones paired with ethanol improved the trap capture compared to traps baited with ethanol alone in southeastern Louisiana. Synergy MTS and FT-II were equally effective in capturing longhorned beetles. The material used to manufacture MTS did not eliminate the need for Fluon application to collect longhorn beetles. Additionally, the location of trapping played a critical role, with distinct habitats supporting unique and diverse groups of cerambycid species. Finally, this study generated novel information on attractants for six species of beetles, namely *O. brunneum*, *P. dentipes*, *L. argentatus*, *L. planidorsus*, *A. haldemani*, and *G. triangulifer*.

Our study provides support for factors to consider when designing trapping experiments to capture cerambycids, especially in subtropical regions. Future studies should aim to identify the specific pheromone components driving responses observed in six beetle species with no previously known attractants, as well as explore subspecies-level differences in pheromone communication. Expanding trapping efforts across additional habitats (e.g., upland hardwoods, mixed stands) and over multiple years would provide valuable insights into the population dynamics and phenology of cerambycid species. In addition, investigating the host-plant associations of species captured in low numbers will help determine whether they represent incidental bycatch or overlooked ecological niches. By integrating these findings into standardized monitoring frameworks, forestry personnel and regulatory bodies can strengthen early detection and rapid response monitoring programs, as well as management of woodboring insects of conservation concern.

## Acknowledgments

We thank Glen Gentry and Will Forbes of the LSU Bob R. Jones Idlewild Research Station, and Joshua Suit and Zack Lemann of Audubon Nature Institute for assistance during site selection and trap installation at field sites. This research was funded in part by US Forest Service (BIL grant 23-DG-11083150-103), Louisiana Department of Agriculture and Forestry, and LSU AgCenter and Department of Entomology. Experiments described herein all comply with current laws of the USA and the authors declare no conflicts of interest.

## Author contributions

CS: Conceptualization; Methodology; Data curation; Formal analysis; Investigation; Visualization; Writing—original draft; Writing—review & editing. TDJ: Conceptualization; Funding acquisition; Methodology; Project administration; Resources; Supervision; Validation; Writing—review & editing. WS: Investigation. EG: Investigation. NJW: Investigation; Data curation. JMS: Investigation; Data curation; Resources. RRS: Resources; Funding acquisition.

