## Supplementary material for "The impact of lure, trap type, and Fluon application on the capture of longhorned beetles (Coleoptera: Cerambycidae) in the subtropical forests of southeastern Louisiana, USA"

**Supplementary materials**

**Table S1**. Generalized linear mixed models evaluated to assess impact of trap features on the capture of longhorned beetles in our study. All models evaluated are presented, with **bolded** models indicating the best model used to determine the impact of trap features on beetle capture. AIC and BIC values are provided. Symbols included in our models indicate additive (+), interactive (*) main effects, and 1/factor indicates random effects. Models with “*” specified include both interactive and additive effects for both factors connected by the asterisk.

| Treatment tested | Full or reduced models used | AIC | BIC |
| --- | --- | --- | --- |
| Trap type | **M1:** **Trap type + lure + 1/trap check date + 1/site**  M2: Trap type * lure + 1/trap check date + 1/site | **1464.1**  1466.0 | **1484.3**  1489.5 |
| Fluon treatment | **Fluon treatment + 1/trap check date + 1/site** | **875.0** | **889.3** |
| Lure type | **M1:** **Trap type + lure + 1/trap check date + 1/site**  M2: Trap type * lure +1/trap check date + 1/site | **1401.1**  1400.9 | **1421.4**  1424.6 |

**Table S2.** Cerambycid species that were used for statistical analysis with their unique days of being captured and their corresponding total.

| **Species** | **# of unique days of beetles captured** | **# of beetles** |
| --- | --- | --- |
| *S. biustus* (LeConte) | 36 | 3422 |
| *N. mucronatus* (Fabricius) | 35 | 465 |
| *C. dentatus* (Newman) | 31 | 197 |
| *E. mucronatum* (Say) | 29 | 193 |
| *X. colonus* (Fabricius) | 31 | 158 |
| *L. angulatus* (LeConte) | 24 | 145 |
| *G. fasciatus* (DeGeer) | 20 | 106 |
| *S. b. minuens* (LeConte) | 21 | 80 |
| *L. argentatus* (Jacquelin du Val) | 19 | 70 |
| *A. haldemani* (LeConte) | 24 | 52 |
| *A. modestus* (Gyllenhal) | 18 | 48 |
| *P. dentipes* (Olivier) | 14 | 44 |
| *G. triangulifer* (Haldeman) | 24 | 40 |
| *O. brunneum* (Forster) | 14 | 40 |
| *A. quadrigibbus* (Say) | 20 | 39 |
| *N. acuminatus* (Fabricius) | 18 | 32 |
| *L. alpha* (Say) | 10 | 27 |
| *L. planidorsus* (LeConte) | 12 | 25 |
| *N. scutellaris* (Olivier) | 16 | 23 |
| *L. transversus* (Gyllenhal) | 4 | 18 |

**Table S3.** Taxonomy and raw numbers of cerambycid beetles that were captured according to species and site in our study in southeastern Louisiana.

| **Subfamily, tribe, and species** | **Audubon** | **Idlewild** | **Total** |
| --- | --- | --- | --- |
| **Disteniinae** |  |  |  |
| **Disteniini** |  |  |  |
| *Distenia undata* (Fabricius) | 0 | 2 | 2 |
| **Prioninae** |  |  |  |
| **Macrotomini** |  |  |  |
| *Mallodon dasytomus* (Say) | 0 | 2 | 2 |
| **Prionini** |  |  |  |
| *Orthosoma brunneum* (Forster) | 0 | 40 | 40 |
| **Cerambycinae** |  |  |  |
| **Bothriospilini** |  |  |  |
| *Knulliana cincta* (Drury) | 0 | 1 | 1 |
| **Clytini** |  |  |  |
| *Neoclytus acuminatus* (Fabricius) | 11 | 21 | 32 |
| (table cont’d.) |  |  |  |
| **Subfamily, tribe, and species** | **Audubon** | **Idlewild** | **Total** |
| *Neoclytus mucronatus* (Fabricius) | 328 | 137 | 465 |
| *Neoclytus scutellaris* (Olivier) | 1 | 22 | 23 |
| *Xylotrechus colonus* (Fabricius) | 15 | 143 | 158 |
| **Curiini** |  |  |  |
| *Curius dentatus* (Newman) | 154 | 43 | 197 |
| **Eburiini** |  |  |  |
| *Eburia quadrigeminata* (Say) | 1 | 2 | 3 |
| **Elaphidiini** |  |  |  |
| *Elaphidion mucronatum* (Say) | 98 | 95 | 193 |
| *Elaphidion tectum* (LeConte) | 1 | 0 | 1 |
| *Enaphalodes atomarius* (Drury) | 0 | 1 | 1 |
| *Parelaphidion aspersum* (Haldeman) | 2 | 0 | 2 |
| **Plectromerini** |  |  |  |
| *Plectromerus dentipes* (Olivier) | 41 | 3 | 44 |
| (table cont’d.) |  |  |  |
| **Subfamily, tribe, and species** | **Audubon** | **Idlewild** | **Total** |
| **Smodicini** |  |  |  |
| *Smodicum cucujiforme* (Say) | 0 | 1 | 1 |
| **Lamiinae** |  |  |  |
| **Acanthocinini** |  |  |  |
| *Acanthocinus obsoletus* (Olivier) | 2 | 0 | 2 |
| *Astylidius parvus* (LeConte) | 4 | 0 | 4 |
| *Astylopsis collaris* (Haldeman) | 0 | 3 | 3 |
| *Astylopsis fascipennis* (Schiefer) | 1 | 4 | 5 |
| *Astylopsis macula* (Say) | 2 | 1 | 3 |
| *Astylopsis perplexa* (Haldeman) | 2 | 0 | 2 |
| *Atrypanius haldemani* (LeConte) | 44 | 8 | 52 |
| *Graphisurus despectus* (LeConte) | 0 | 2 | 2 |
| *Graphisurus fasciatus* (DeGeer) | 7 | 99 | 106 |
| *Graphisurus triangulifer* (Haldeman) | 14 | 26 | 40 |
| (table cont’d.) |  |  |  |
| **Subfamily, tribe, and species** | **Audubon** | **Idlewild** | **Total** |
| *Hyperplatys aspersa* (Say) | 0 | 1 | 1 |
| *Leptostylopsis argentatus* (Jacquelin du Val) | 38 | 32 | 70 |
| *Leptostylopsis planidorsus* (LeConte) | 18 | 7 | 25 |
| *Leptostylus asperatus* (Haldeman) | 0 | 9 | 9 |
| *Leptostylus transversus* (Gyllenhal) | 9 | 9 | 18 |
| *Lepturges angulatus* (LeConte) | 110 | 35 | 145 |
| *Lepturges confluens* (Haldeman) | 1 | 2 | 3 |
| *Liopinus alpha* (Say) | 13 | 14 | 27 |
| *Styloleptus biustus* (LeConte) | 2067 | 1355 | 3422 |
| *Styloleptus b. minuens* (LeConte) | 71 | 9 | 80 |
| **Acanthoderini** |  |  |  |
| *Aegomorphus modestus* (Gyllenhal) | 3 | 45 | 48 |
| *Aegomorphus morrisii* (Uhler) | 0 | 1 | 1 |
| *Aegomorphus quadrigibbus* (Say) | 26 | 13 | 39 |
| **Subfamily, tribe, and species** | **Audubon** | **Idlewild** | **Total** |
| **Dorcaschematini** |  |  |  |
| *Dorcaschema alternatum* (Say) | 12 | 0 | 12 |
| **Pogonocherini** |  |  |  |
| *Ecyrus dasycerus* (Say) | 12 | 2 | 14 |
| *Pogonocherus mixtus* (Haldeman) | 0 | 1 | 1 |
| **Pteropliini** |  |  |  |
| *Ataxia crypta* (Say) | 2 | 0 | 2 |
| *Ataxia falli* (Breuning) | 2 | 0 | 2 |
| **Grand total** | **3112** | **2191** | **5303** |

**Table S4**. Cerambycid species that were significantly impacted by site with their estimate, SE, and p-value in southeastern Louisiana. Note: this table was generated by treating site as fixed effect in the GLMM model.

| **Species** | **Audubon** | **Idlewild** | **Total** | **Estimate ± SE** | **z-value** | **p-value** |
| --- | --- | --- | --- | --- | --- | --- |
| *A. modestus* | 3 | 45 | 48 | 2.66 ± 0.61 | 4.34 | <0.01 |
| *A. quadrigibbus* | 26 | 13 | 39 | -0.80 ± 0.38 | -2.12 | 0.04 |
| *A.* haldemani | 44 | 8 | 52 | -1.84 ± 0.40 | -4.56 | <0.01 |
| *C. dentatus* | 154 | 43 | 197 | -1.30 ± 0.21 | -6.16 | <0.01 |
| *E. dasycerus* | 12 | 2 | 14 | -1.71 ± 0.84 | -2.02 | 0.04 |
| *G. fasciatus* | 7 | 99 | 106 | 2.65 ± 0.39 | 6.76 | <0.01 |
| *L. planidorsus* | 18 | 7 | 25 | -0.99 ± 0.49 | -2.02 | 0.04 |
| *L. angulatus* | 110 | 35 | 145 | -1.10 ± 0.27 | -4.03 | <0.01 |
| *N. mucronatus* | 328 | 137 | 465 | -0.94 ± 0.19 | -7.94 | <0.01 |
| *N. scutellaris* | 1 | 22 | 23 | 3.07 ± 1.03 | 2.99 | 0.003 |
| *P. dentipes* | 41 | 3 | 44 | -3.03 ± 0.71 | -4.24 | <0.01 |
| *S. biustus* | 2067 | 1355 | 3422 | -0.43 ± 0.04 | -12.26 | <0.01 |
| *S. b. minuens* | 71 | 9 | 80 | -2.09 ± 0.35 | -5.88 | <0.01 |
| *X. colonus* | 15 | 143 | 158 | 2.24 ± 0.27 | 8.26 | <0.01 |

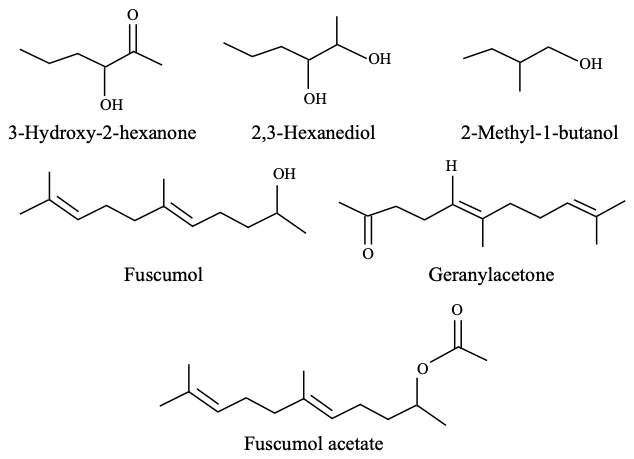

**Figure S1.** Chemical structures of each component present in a blend of cerambycid pheromones lure.

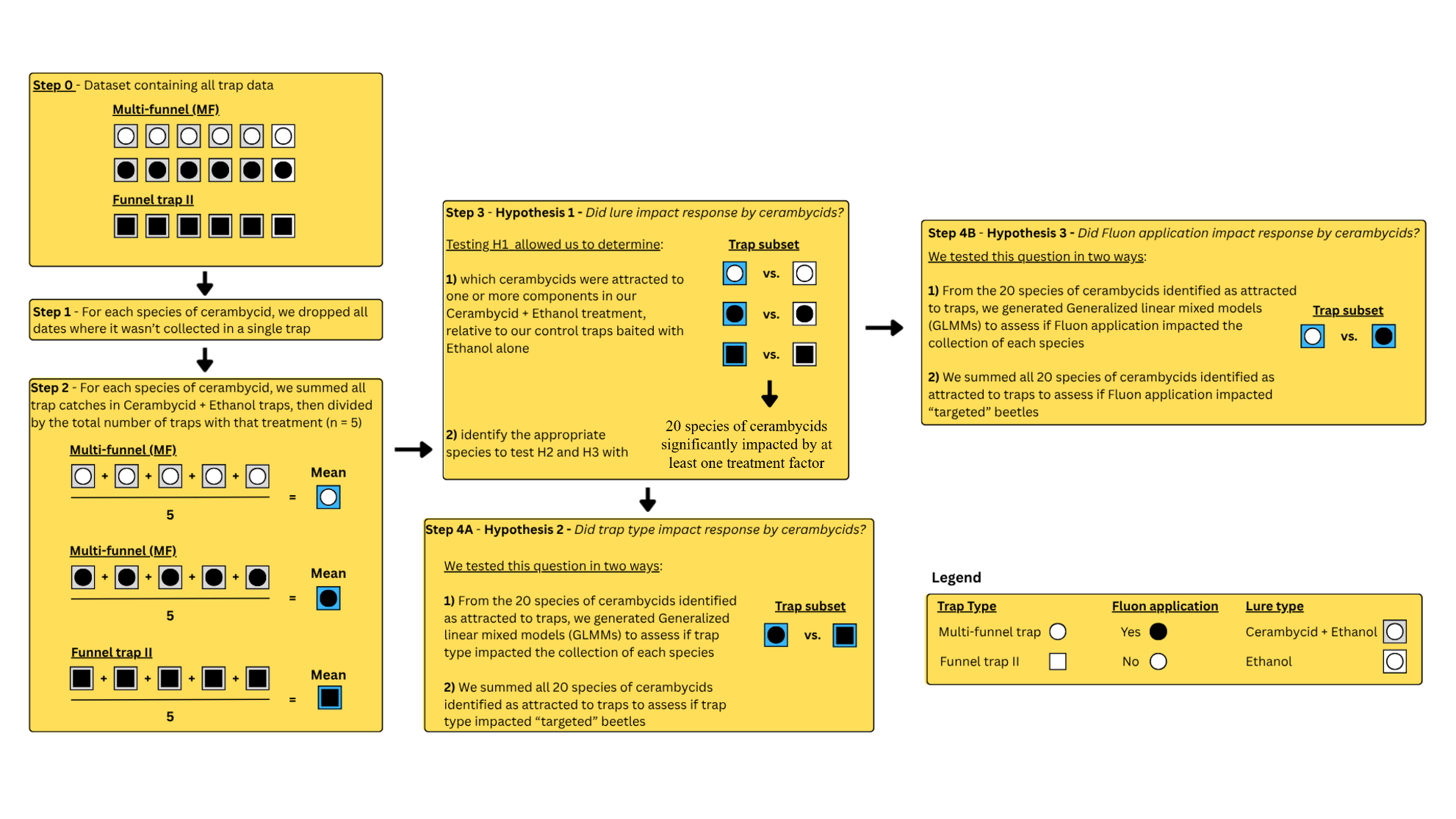
**Figure S2.** Flowchart of data curation to determine the impact of lure (H1), trap type (H2), and Fluon application (H3) on the capture of longhorned beetles in our study in southeastern Louisiana. Each sequential box on the flowchart shows a step in which data were modified or subsetted to facilitate testing our hypotheses. Each box that uses or modifies trap data according to treatment is visualized with different shapes and colors; Multi-funnel traps (MF) are represented by a circle, Funnel trap II (FT-II) are represented by squares; The fill of these shapes indicates whether traps were paired with the cerambycid pheromone blend and low release ethanol lure (gray’ C+E), or a low release ethanol lure (white; E); all circles and squares have a second shape of the same type inside of it, which represents whether it received a Fluon application (black), or no Fluon application (white). Note that FT-II only received Fluon treatment. In step 2, the mean was calculated for all MF and F-II with the C+E lure blend to facilitate analysis. The mean of these five traps per treatment are represented by a blue fill rather than gray.
